# Modular small RNA drives the emergence of virulence traits and environmental trade-offs in *Vibrio cholerae*

**DOI:** 10.1101/2024.07.30.605856

**Authors:** Deepak Balasubramanian, Cole Crist, Salvador Almagro-Moreno

## Abstract

The sole gain of laterally acquired virulence genes does not fully explain the transition of environmental strains into human pathogens. To date, the specific molecular drivers and fitness trade-offs that enable some strains within a population to undergo this process remain enigmatic. Here, we describe a small RNA (sRNA) with a unique modular structure that shapes the evolution of toxigenic *Vibrio cholerae*, the agent of cholera. The sRNA comprises of a highly variable 5’ module located within the *ompU* ORF and a conserved 3’ one downstream from the gene. This atypical location confers a distinct bimodular structure to the OmpU-encoded sRNA (OueS), generating allelic variants that differentially contribute to the emergence of virulence potential in some strains and associated fitness trade-offs between human infection and environmental survival. Unlike environmental counterparts, the OueS allele from toxigenic strains controls phenotypes essential during host colonization: a) stringently inhibits biofilm formation via a novel sRNA-sRNA interaction by suppressing the iron-responsive sRNA RyhB, and b) confers resistance against intestinal bacteriophages by activating the recently discovered CBASS phage defense system. Toxigenic OueS is also required for successful intestinal colonization and, as the first known example of its kind, acts as a functional surrogate of the master virulence regulator ToxR, controlling over 84% of its regulome. On the other hand, isogenic strains encoding environmental alleles of OueS exhibit higher competitive fitness than those harboring toxigenic ones during colonization of natural reservoirs such as crustaceans and phytoplankton. Our findings provide critical insights into the evolution of toxigenic *V. cholerae* and reveal specific molecular mechanisms and fitness costs associated with the emergence of pathogenic traits in bacteria.

## INTRODUCTION

The emergence of human pathogens is one of the most pressing public health concerns of modern times. Pathogen emergence is a complex and multifactorial phenomenon that culminates with the organism acquiring the ability to colonize and harm the human host^1–3^. A successful infectious process necessitates accomplishing numerous steps and overcoming critical host-associated hurdles. These include avoidance of host defense mechanisms and phage predation, the correct spatiotemporal expression of virulence genes in response to host signals, and, ultimately, proliferation and transmission. Despite its critical relevance, many essential questions regarding this complex phenomenon remain unanswered. For instance, the sole gain of laterally acquired virulence genes cannot fully explain the transition of environmental strains into human pathogens, as typically only a limited number of clades within a species eventually emerge to cause disease. To date, the specific molecular drivers and fitness trade-offs that enable certain strains within a population to undergo this process remain enigmatic. Identifying the elements associated with the process of pathogen emergence is critical for successful disease management and control^4^.

*Vibrio cholerae* is the etiological agent of the severe diarrheal disease cholera and remains a major scourge in places with limited access to clean drinking water^5,6^. The disease is contracted via the consumption of food or water contaminated with toxigenic strains of *V. cholerae*^5^. Following ingestion in the form of planktonic cells or forming biofilm, *V. cholerae* must overcome several host defenses such as acidic stomach pH, bile acids, and antimicrobial peptides^7–10^. Successful colonization also requires stringent downregulation of biofilm formation, as it triggers a strong immune response, paired with upregulation of flagella-mediated motility to swim through the mucus layer towards the epithelium^11–13^. A complex regulatory cascade mediated by the virulence regulator ToxR facilitates adaptation to the harsh conditions within the host and, along with other regulators, the coordinated and timely expression of virulence genes^14–17^. ToxR is a transmembrane protein with a cytoplasmic DNA-binding domain that controls the expression of hundreds of genes, however, the regulator only binds directly to the promoter of 39 genes^15,18–21^. Among the genes directly regulated by ToxR are those encoding the cholera toxin (CT), the source of the watery diarrhea associated with cholera, and, together with TcpP, the expression of ToxT, a master virulence regulator^18,22^. ToxR also directly and reciprocally modulates the expression of two outer membrane proteins (OMPs): OmpU and OmpT. Specifically, ToxR represses *ompT* expression whereas it activates transcription of *ompU*^19,23,24^.

OmpU is highly expressed during infection, making up to 60% of the outer membrane, and is essential for host survival and intestinal colonization^19,25,26^. The porin is associated with numerous virulence-adaptive phenotypes such as bile resistance, or tolerance to organic acids and cationic peptides (e.g. polymyxin B or the host-derived antimicrobial peptide P2)^7,8,19,26,27^. Beyond resistance to cell damaging agents, OmpU has also been associated with several more complex phenotypes. For instance, OmpU exerts immunomodulatory effects such as induction of host tolerance to LPS and suppression of the host immune response by inducing M1 polarization of macrophages^28,29^. Furthermore, OmpU serves as the receptor of bacteriophage ICP2, a phage isolated from stool samples of cholera patients^30^. We recently determined that OmpU is also associated with biofilm formation in *V. cholerae*, as *ompU* deletion mutants form 3-4-fold more robust biofilms than wild-type strains^31^. Interestingly, moonlighting roles critical to infection-related processes appear to be a common feature among OMPs in bacterial pathogens. For instance, OMPs have been associated with activation of host signal transduction pathways, inhibition of host cell apoptosis, adhesion and signaling, and modulation of immune responses^32–36^. To date, the mechanisms by which porins like OmpU modulate such diverse virulence-associated phenotypes remain poorly understood.

Despite encompassing over 200 serogroups, only *V. cholerae* strains from serogroups O1 and O139 can cause cholera in humans^37–39^. Strains from these two serogroups are phylogenetically related and confined to a single clade containing all toxigenic *V. cholerae* called the pandemic cholera group (PCG)^40^. To elucidate the evolutionary origins and emergence of toxigenic *V. cholerae*, we previously developed a comparative genomics framework^31^. Our analyses of environmental and toxigenic strains of *V. cholerae* identified the presence of allelic variations in core genes that circulate in environmental populations and confer preadaptations to host colonization, enhancing their pandemic potential^31^. These genes encode what we term virulence adaptive polymorphisms (VAPs) and are connected with the emergence of toxigenic strains of *V. cholerae*, explaining their limited phylogenetic distribution^31^. Interestingly, we found that *ompU* in toxigenic *V. cholerae* strains encode VAPs and the virulence traits associated with the porin are allelic-dependent, as these traits are absent from strains with *ompU* alleles that do not encode them^31,41^. The molecular mechanisms controlling these allelic-dependent emergent phenotypes remain unaddressed.

Here, we describe a small RNA (sRNA) with a unique modular structure that shapes the evolution of toxigenic strains of *Vibrio cholerae*, the agent of cholera. The sRNA comprises of a highly variable 5’ module located within the *ompU* ORF and a conserved 3’ one downstream from the gene. The bimodular nature of the OmpU-encoded sRNA (OueS) generates allelic variants that differentially contribute to **a)** the emergence of virulence potential in toxigenic strains, and **b)** associated fitness trade-offs between human infection and environmental survival. Our findings provide critical insights into the evolution of pandemic *V. cholerae* and reveals specific molecular mechanisms and fitness costs associated with the emergence of pathogenic traits in bacteria.

## RESULTS

### OmpU represses biofilm formation in *V. cholerae* and modulates its transcriptome

OmpU plays a major role in *V. cholerae* pathogenesis, nonetheless, it remains unclear how an outer membrane porin can modulate so many disparate virulence-related phenotypes^7,8,19,26,27^. We previously demonstrated that deletion of *ompU* in toxigenic *V. cholerae* leads to more robust biofilm formation^31^. To dissect this phenotype, we ectopically expressed *ompU* from an inducible plasmid in an in-frame isogenic *ompU* deletion mutant in *V. cholerae* C6706, an El Tor strain from the seventh cholera pandemic (Δ*ompU*-pOmpU^C6706^). Expression of *ompU* decreases biofilm formation in the Δ*ompU* mutant back to wild-type (WT) levels, indicating that OmpU represses biofilm formation in *V. cholerae* (**Fig. 1A**). On the other hand, expression of an environmental *ompU* allele from a strain isolated from the Great Bay Estuary in New Hampshire, *V. cholerae* GBE1114^42^, either ectopically or through allelic exchange in the isogenic C6706 background, does not repress biofilm formation, suggesting this phenotype is allelic-dependent (**Fig. 1A**)^31^. Interestingly, biofilm suppression is critical in the early stages of intestinal colonization as biofilm components elicit a strong immune response leading to clearance of *V. cholerae* strains that do not timely repress it^43^. We found it quite puzzling that expression of a porin localized in the outer membrane can control a phenotype that is central to the infectious process.

**Figure 1.**
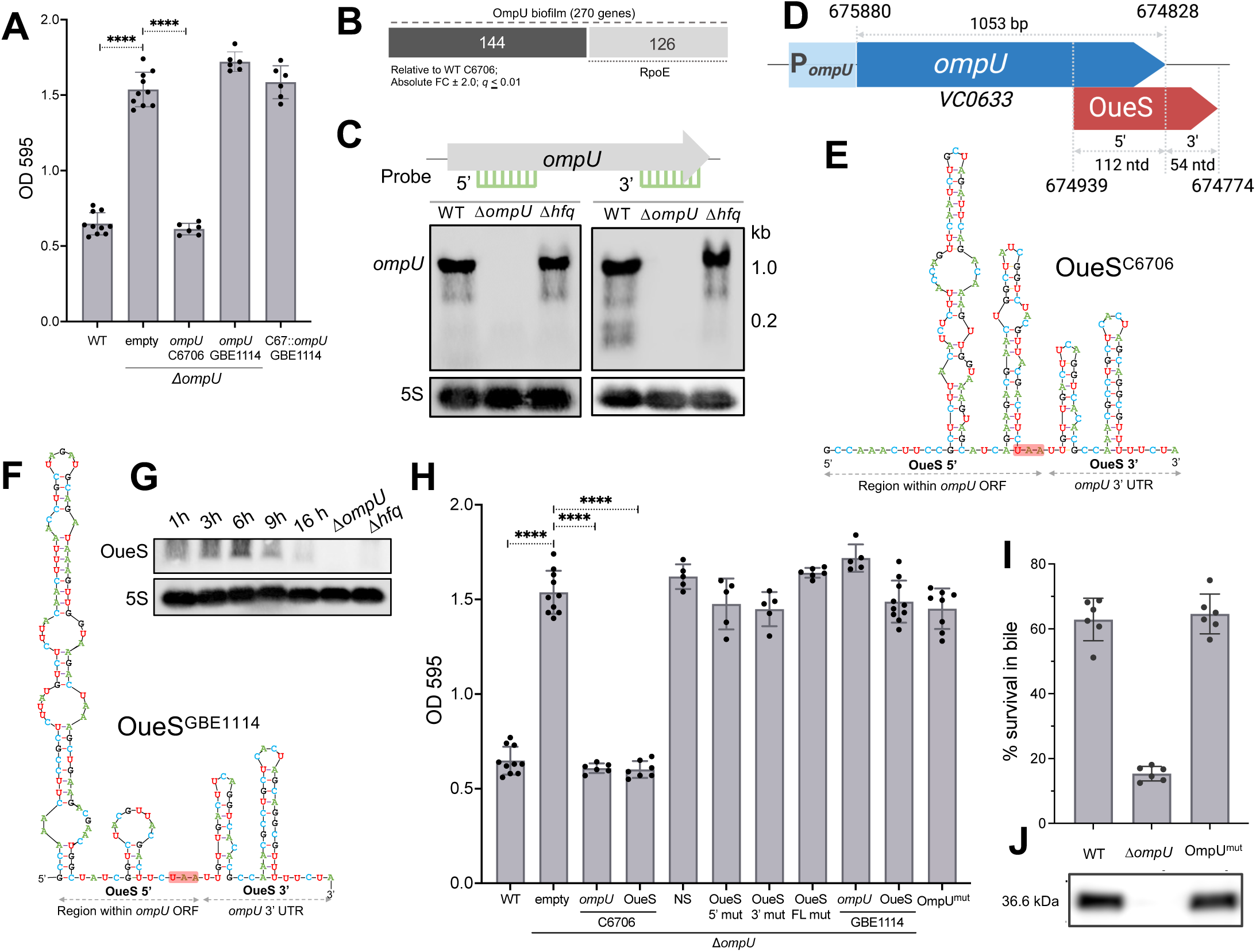
The *ompU* ORF encodes a small RNA with a unique modular structure. (A) Biofilm formation of *V. cholerae* C6706 WT, Δ*ompU* and Δ*ompU* strains expressing OmpU^C6706^ or OmpU^GBE1114^, either in *cis* or *trans*. Strains harbor either the empty vector or the *ompU*-encoding construct. Strains expressing OmpU from toxigenic strains repress biofilm formation. n ≥ 6. Two-tailed T-test *p*-value **** ≤ 0.0001. (B) Total number of differentially regulated genes (absolute FC ≥ 2.0; *q* ≤ 0.01) from the transcriptomes of *V. cholerae* C6706 were compared against Δ*ompU* and Δ*rpoE* strains isolated from biofilms. (C) Total RNA was isolated from *V. cholerae* C6706 WT, Δ*ompU* and Δ*hfq* cells and subjected to northern blot analyses using probes specific for the 5’ and 3’ ends of the *ompU* ORF. The probes specific for the 3’ end detect additional bands in the 150-200 ntd region in the WT strain that are absent in the *ompU* and *hfq* deletion mutants. 5S rRNA-specific probes were used as a loading control. (D) Schematic of the genomic location, coordinates, and termini of the newly identified *ompU*-encoded sRNA (OueS) based on 5’ and 3’ RACE data. Modular structural variability is seen in the predicted secondary structures of OueS from (E) toxigenic (C6706) and (F) environmental (GBE1114) *V. cholerae* strains. In E and F, the 5’ region of OueS encoded within the *ompU* ORF and the 3’ region that lies within the genes UTR are indicated. The stop codon of the *ompU* ORF is highlighted in red. (G) Expression of OueS was examined by northern blotting using total RNA isolated at various stages of growth from WT *V. cholerae* C6706. OueS expression peaks during the early stationary growth phase (6h). (H) Expression of OueS from toxigenic strain *V. cholerae* C6706 suppresses biofilm formation. Strains harbor either the empty vector or encode variants of *ompU* or OueS. n ≥ 5. Wobble codon-based mutations of the genomic OueS locus (OmpU^mut^) does not affect (I) OmpU porin function as determined by resistance to 0.4% whole bile, or (J) expression and stability as seen in OmpU immunoblots.

To gain insights into the mechanisms behind this marked phenotype, we examined the transcriptional changes associated with OmpU-mediated biofilm repression. We performed RNA-Seq analyses of *V. cholerae* biofilms produced by WT, Δ*ompU* and Δ*ompU*-pOmpU^C6706^ strains. Our data reveals a drastic shift in the transcriptome of the Δ*ompU* mutant strain compared to WT (**Fig S1**), with differential regulation of 270 genes of which 181 genes are repressed by OmpU and 89 are activated (absolute FC ≥ 2.0, *q* ≤ 0.01) (**Fig 1B; Table S1**). Ectopic expression of *ompU*^C6706^ in the Δ*ompU* background results in a shift of the transcriptome, relatively closer to WT, and the recovery in expression of the dysregulated genes (**Fig S1; Table S1**). GO enrichment analysis reveals that OmpU primarily represses genes involved in iron metabolism and acquisition, specifically in vibriobactin metabolism, siderophore transport, and iron ion homeostasis (*q* ≤ 0.05) (**Fig S1; Table S1**). On the other hand, OmpU activates genes involved in the defense response mechanisms to phages and symbionts, as well as those involved in heat response and protein folding (*q* ≤ 0.05) (**Fig S1; Table S1**).

Upon misfolding, OmpU activates the cell envelope stress response via the release of sigma factor RpoE in the cytosol^44^. To address the potential role of RpoE in the transcriptomic changes associated with OmpU, we examined the biofilm transcriptome of a Δ*rpoE* mutant strain (**Fig 1B**). Comparison of the transcriptome datasets of Δ*ompU* and Δ*rpoE* strains, relative to WT, demonstrates that although there is a natural overlap between the two datasets, a large percentage of the transcriptomic changes associated with *ompU* expression during biofilm formation appear to be independent of RpoE activation (**Fig 1B; Table S2**). Specifically, OmpU modulates the expression of 270 genes during biofilm formation and 144 of those genes (53.3%) are not associated with RpoE (**Fig 1B; Table S2**). Overall, our data indicates that OmpU represses biofilm formation in an allele-dependent manner and has a significant impact on the *V. cholerae* transcriptome.

### The *ompU* ORF encodes a modular small RNA

Given the large transcriptomic changes associated with OmpU extending beyond the activation of RpoE, we considered the possibility of the *ompU* gene coding for regulatory sequences within its ORF besides the porin. A genome-wide screening in *V. cholerae* indicated, among thousands of others, potential sRNAs encoded within the 3’ end of the *ompU* gene^45^. To examine whether the *ompU* ORF potentially encodes an internal sRNA, we performed northern blot analysis with probes targeting the 3’ termini of *ompU* and the 5’ termini as a control. We reasoned that a probe targeting the sRNA locus within *ompU* should result in two bands corresponding to both the *ompU* mRNA and the potential sRNA. The northern blot data shows that a probe specific for the 5’ end of the *ompU* ORF (675,493–675,527 ntd) results in a ∼1100 ntd band corresponding to the expected size of the mRNA transcript (**Fig 1C**). On the other hand, in addition to the ∼1.1 kb mRNA band, smaller size bands of ∼200 ntds appear with a probe targeting the 3’ end of the *ompU* ORF (674,837–674,876 ntd) potentially corresponding to a sRNA (∼160 ntd) (**Fig 1C**). Hfq is an sRNA binding protein that plays an integral role in their stability and target recognition^46^. Interestingly, those lower bands are not detectable in a Δ*hfq* mutant, strengthening the possibility of a sRNA being encoded within the 3’ end of the *ompU* ORF (**Fig 1C**).

To define the 5’ and 3’ termini of the potential sRNA encoded within the *ompU* ORF, we performed rapid amplification of cDNA ends (RACE) assays. Given the overabundance of the *ompU* transcript and the overlapping nature of the mRNA and the potential sRNA, our initial 5’ RACE studies yielded only the termini of ∼1100 bp *ompU* mRNA transcript. To circumvent this, total RNA from the WT strain cultured in LB was size fractionated on a MOPS agarose gel and the 100-500 ntd fraction, encompassing the potential sRNA region was extracted and used as template for RACE studies. Assays on this size-fractionated RNA led to the identification of a sRNA corresponding to 166 ntds that we termed *<u>o</u>mp*U-<u>e</u>ncoded <u>s</u>mall RNA (OueS) (**Fig 1D**). Surprisingly, even though the majority of the sRNA (112 ntds) is encoded within the *ompU* ORF before its 3’ terminus, 54 ntds are located downstream of the stop codon in the 3’ UTR (674,939 – 674,774 on chromosome I), making the OueS locus quite atypical as it overlaps an ORF and its 3’ UTR (**Fig 1D**). Computational prediction of the secondary structure of OueS reveals a four stem-loop structure, with the terminal stem-loop ending in a poly-U tail typical of a Rho-independent terminator and a potential Hfq binding site, further corroborating our finding in Fig 1C^46,47^ (**Fig 1E**). Interestingly, the environmental *ompU* allele GBE1114, which does not lead to biofilm suppression, encodes a distinct structural variant of OueS (**Fig 1F**). Even though the length of the toxigenic and environmental OueS alleles is 166 ntds, their predicted secondary structures drastically differ in the 5’ region encoded within the *ompU* ORF but is identical in the OueS region downstream of the *ompU* gene that harbors the Hfq binding motif^47^, suggesting a bimodular structure of OueS (**Fig 1E, 1F**). The non-canonical location of OueS, between an ORF and its UTR, and its modular nature makes this a unique sRNA that might contribute to the phenotypic diversity of *V. cholerae*^48–50^. Monitoring OueS expression at different stages of growth by northern blotting indicate that its expression peaks at the early stationary phase (6h) with a reduction in sRNA levels at later growth phases (**Fig 1G**). Overall, our results indicate that the *ompU* ORF in *V. cholerae* encodes both the OmpU porin and the 5’ terminus of a modular 166 ntd sRNA that we termed OueS.

### OueS represses biofilm formation in *V. cholerae*

Next, we examined whether the *ompU*-mediated biofilm repression seen in **Fig 1A** is mediated by OueS. We ectopically expressed OueS in the Δ*ompU* mutant (Δ*ompU*-pOueS) and determined that the strain recovers the WT phenotype and displays reduced biofilm formation when compared to Δ*ompU*, similar to the ectopic expression of OmpU (**Fig 1H**). This finding indicates that OueS is responsible for the *ompU*-mediated biofilm repression in *V. cholerae* (**Fig 1H**). To examine the specificity of OueS-mediated biofilm repression, first we ectopically expressed a non-specific (NS) region outside of the OueS locus located towards the 5’ end of the *ompU* ORF (675,307–675,521). The NS region had no effect on biofilm formation compared to the Δ*ompU* strain (**Fig 1H**). Subsequently, we planned on generating mutations within OueS to potentially nullify its effect. However, the 5’ end of OueS is encoded within the *ompU* ORF and thus, mutations affecting OueS would also affect the OmpU coding region. To ensure that the mutations within the sRNA do not disrupt the protein coding sequence, we took advantage of the codon wobble and mutated every third base in the 166 bp OueS locus, disrupting the sRNA secondary structure while preserving the OmpU amino acid sequence (**Fig S2**), what we termed OueS full-length mutant (FL-mut). We integrated OueS FL-mut at the native OueS locus generating a strain encoding a WT OmpU porin and a mutated OueS sRNA (C6706 OmpU^mut^). The OmpU porin produced by this strain is functional and stable, as determined by resistance to bile (**Fig 1I**) and OmpU immunoblots (**Fig 1J**), respectively, indicating that the porin function remains unaffected due to OueS disruption. The OmpU^mut^ strain formed higher biofilm levels, similar to Δ*ompU*, further indicating that biofilm repression is mediated by OueS and not OmpU (**Fig 1H**). In addition, we generated ectopic expression constructs of three OueS wobble mutants: OueS 5’-mut harboring mutations exclusively in the OueS region that lies within the *ompU* ORF, and OueS 3’-mut with mutations only in the OueS region downstream of *ompU*, and the OueS FL-mut with mutations across the entire length of the sRNA. All three mutant versions of OueS disrupted the predicted secondary structures of the sRNA (**Fig S2**). Ectopic expression of either of the three OueS mutant sRNAs (5’-mut, 3’-mut and FL-mut) in the Δ*ompU* strain does not lead to suppression of biofilm formation (**Fig 1H**). Additionally, unlike the OmpU and OueS alleles from toxigenic strain C6706, ectopic expression of the environmental OueS^GBE1114^ does not lead to biofilm suppression (**Fig 1H**). Overall, our results demonstrate that **a)** OueS leads to stringent biofilm repression in *V. cholerae*, **b)** this phenotype is dependent on the OueS allele and **c)** the sRNA has a unique bimodular structure that divides its *ompU*-encoded 5’ end from its 3’ terminus.

### OueS is responsible for *ompU-*dependent transcriptomic changes during biofilm formation

OmpU modifies the transcriptome of *V. cholerae* during biofilm formation (**Fig 1B** and **Fig S1**). Given the effect of OueS on biofilm repression, we hypothesized that expression of OueS would be responsible for this altered transcriptional profile. To address this, we examined the transcriptomes of Δ*ompU*-pOueS^C6706^ cells harvested from biofilms. Comparison of the biofilm transcriptomes of Δ*ompU*-pOueS^C6706^ to those of the WT and Δ*ompU* strains reveal that OueS expression restores the shift in the transcriptome similar to ectopic *ompU* expression (**Fig S1A**). Further, we compared the differentially regulated genes (absolute FC ≥ 2.0, *q* ≤ 0.01) of two pairwise comparisons: WT Vs Δ*ompU* (270 genes) and Δ*ompU*-pOueS^C6706^ Vs Δ*ompU* (267 genes). There are 132 genes (48.89%) in the overlap region whose expression is modulated by OueS, 101 of which are repressed and 31 activated (**Table S3)**. Analysis of the expression values of these 132 genes between the two pair-wise comparisons reveals that the direction of the fold change (activation Vs repression) as well as the expression levels are very similar in 131 of the 132 genes between the two groups (Pearson Correlation Coefficient = 0.9998; **Table S3**). GO enrichment analysis on the genes repressed by OueS reveal that 63 of the 101 OueS repressed genes are enriched in the GO Biological Process database and 17 of the 31 activated ones are enriched. Similar to the OmpU transcriptome, among the OueS-repressed genes there was a significant positive enrichment (*q* < 0.05) of genes belonging to the vibriobactin biosynthetic processes and siderophore transmembrane transport (**Fig 2A**). Overall, our data indicates that OueS is responsible for changes in the transcriptome during biofilm formation associated with OmpU.

**Figure 2.**
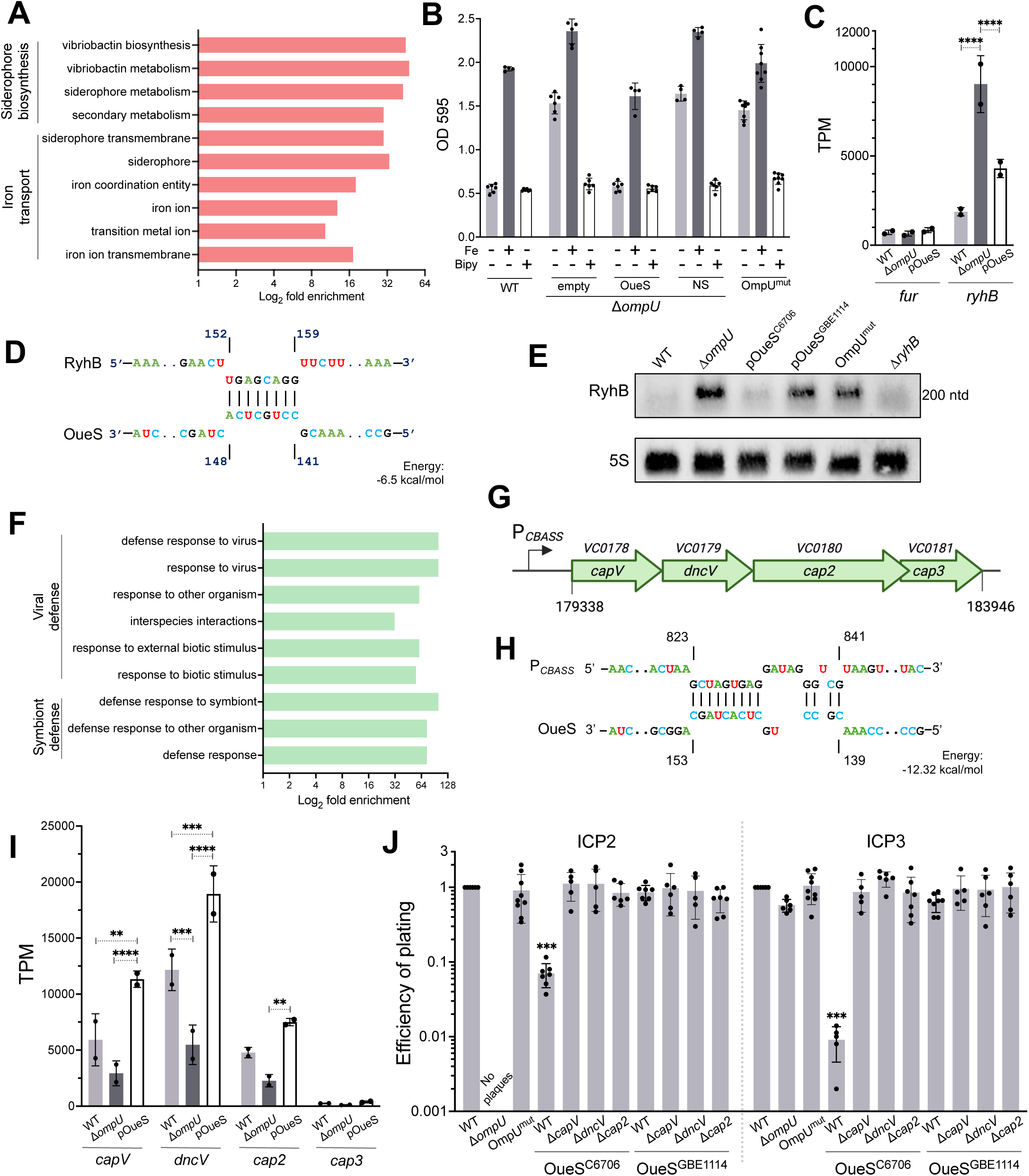
OueS controls phenotypes essential for intestinal colonization. Biofilm repression is critical for successful intestinal colonization as one biofilm component, the mannose sensitive hemagglutinin, elicits a prohibitively strong immune response^43^. (A) Functional gene ontology (GO Biological Processes) categorization of the genes repressed by OueS during biofilm formation was performed by the Panther Overrepresentation Test^88,89^. OueS represses categories related to iron uptake (B) Biofilm formation of strains expressing mutant versions of OueS were examined under iron-replete (dark grey bars) and iron-depleted (white bars) conditions by exogenous addition of 100 µM FeCl_3_ or 100 µM 2,2’-bipyridyl, respectively. Data was compared to typical growth media for baseline iron conditions (light grey bars). (C) Normalized read counts (transcripts per kilobase million, TPM) from the biofilm transcriptome data indicates OueS-mediated repression of *ryhB* expression. TPM data was normalized to expression of *VC0799* whose expression remains stable between the strains (TPM range 18 – 33). (C) Sequence homology predicts interaction of OueS with RyhB. (E) Northern blots demonstrate increased levels of RyhB in a Δ*ompU* mutant. The presence of RyhB is drastically reduced upon expression of OueS^C6706^. Resistance to lytic bacteriophages contributes to intestinal survival, host colonization, and seasonality of cholera outbreaks ^55^. (F) Functional GO analysis^88,89^ of OueS activated genes reveals enrichment of viral and symbiont defense categories. (G) Schematic of the cyclic oligonucleotide-based anti-phage signaling system (CBASS) operon: *capV* (VC0178)*, dncV* (VC0179)*, cap2* (VC0180), and *cap3* (VC0181). The genomic coordinates pertaining to *V. cholerae* El Tor N16961 genome are indicated. (H) Sequence homology predicts interaction of OueS with the promoter of the CBASS operon. (I) TPM data from the transcriptome of biofilm-derived cells reveals OueS-mediated activation of CBASS operon genes. TPM data normalization was performed as in (C) above. (J) Plaque assays reveal that ectopic expression of the toxigenic OueS^C6706^ allele in WT strain confers resistance to *V. cholerae* bacteriophages ICP2 and ICP3, in contrast to its environmental counterpart OueS^GBE1114^. Deletion of genes in the CBASS operon abolishes the effect of OueS^C6706^ on phage resistance. 2way ANOVA multiple comparisons test *p*-value: ** <0.01, *** <0.001, **** <0.0001. *p*-values are in comparison to the WT, unless specified.

### OueS from toxigenic strains suppresses biofilm formation by repressing the iron-responsive sRNA RyhB

We analyzed the OueS-regulated genes for insights into the molecular basis behind the stringent biofilm repression associated with the toxigenic allele of the sRNA. Iron is important for biofilm formation and OueS^C6706^ downregulates the expression of numerous iron uptake genes^51–53^ (**Fig 2A**; **Table S3**). Consequently, we examined the interplay between iron availability and biofilm formation, and their association with OueS. Addition of exogenous iron to the media results in increased biofilm formation across all the strains, overriding the suppression exerted by OueS (dark grey bars, **Fig 2B**). On the other hand, depletion of the native iron in the media using the iron chelator 2,2’-bipyridyl results in drastically reduced biofilm formation in all strains, closely resembling the phenotype associated with OueS production (white bars, **Fig 2B**). These changes during biofilm formation are not due to growth defects of the strains under these conditions (**Fig S3A-C**). Our data indicates that the transcript numbers of the major iron uptake repressor *fur* (*VC2106*) are not significantly altered by either deletion of *ompU* or ectopic OueS expression (**Fig 2C**)^53^. In contrast, OueS modulates the transcript numbers of *ryhB*, the Fur-regulated sRNA (**Fig 2C**). RyhB enhances biofilm formation by modulating iron uptake^51^, suggesting a mechanistic basis for biofilm regulation via OueS-mediated repression of the sRNA (**Fig 2C**). Computational analysis predicts the interaction of OueS^C6706^ with RyhB based on sequence homology (**Fig 2D**). Using northern blots, we examined the stability of RyhB in different *V. cholerae* OueS mutant strains (**Fig 2E**)^51^. Our results show that strains encoding OueS^C6706^ exhibit undetectable levels of RyhB (**Fig 2E**). This repression is relieved in strains coding for the non-functional OueS (OmpU^mut^) or an environmental allele of OueS (OueS^GBE1114^), corroborating our biofilm data (**Fig 2E**). Overall, our results indicate that the biofilm inhibition meditated by OueS involves suppression of RyhB in an allelic-dependent manner.

### Toxigenic OueS confers resistance against intestinal bacteriophages by activating the CBASS defense system

Our transcriptome analyses reveal that the genes activated by OueS^C6706^ are primarily involved in the ‘defense response to virus’ and ‘defense response to symbionts’ categories, resembling the functional enrichment seen in the OmpU biofilm transcriptome (**Fig 2F, Table S1**). The four-gene operon encoding the cyclic oligonucleotide-based anti-phage signaling system (CBASS) is located on the *Vibrio* Seventh Pandemic Island 1 (VSP-I) (**Fig 2G**)^54^. Computational analysis predicts potential interaction of OueS^C6706^ upstream of the *VC0178* ORF, the region that encompasses the promoter of the operon (**Fig 2H**). Furthermore, transcriptomic analyses indicate that genes in the CBASS operon are regulated in an OueS-dependent manner (**Fig 2I**). Specifically, compared to the WT, the first three genes of the operon (*capV*, *dncV* and *cap2*) exhibit a decrease in expression in the Δ*ompU* strain (**Fig 2I**, **Table S3**). Expression levels of these genes drastically increase upon expression of OueS in the Δ*ompU* background reverting or exceeding WT levels of expression (**Fig 2I**).

Given the critical role of bacteriophages in shaping *V. cholerae* intestinal infection and cholera outbreak dynamics^30,55–57^, we examined the role of OueS in modulating sensitivity to ICP1, ICP2 and ICP3 phages^30,58,59^. These phages either utilize the OmpU porin (ICP2) or O1 antigen of LPS (ICP1 and ICP3) as receptors^30,58,59^. Phage ICP1 generated no plaques in any of the strains tested, possibly due to phase variations in the poly(A) tract of the *wbeL* gene in *V. cholerae* C6706 resulting in altered sensitivity^60^. On the other hand, plaque assays reveal that toxigenic OueS confers resistance to ICP2 and ICP3 (**Fig 2J**). As previously determined, the Δ*ompU* strain is resistant to ICP2 infection due to absence of its OmpU receptor^30,59^, and shows a moderate reduction in ICP3 plaquing efficiency (**Fig 2J**). Since the presence of OmpU affects resistance to both ICP2 and ICP3, and OueS exhibits strong temporal and condition-dependent expression, for plaque assays we examined the role of OueS in phage resistance by ectopic expression of toxigenic OueS (OueS^C6706^) and its environmental counterpart (OueS^GBE1114^) (**Fig. 2J**). WT cells expressing OueS^C6706^ show a drastic reduction in plaquing efficiency for both ICP2 (∼1-log) and ICP3 (∼2-log) (**Fig 2J**). Conversely, strains ectopically expressing OueS^GBE1114^ exhibit a similar plaquing efficiency as the WT strain, indicating allelic dependency of this phenomenon. Subsequently, we examined the role of CBASS components in this phenotype. Expression of OueS^C6706^ in deletion mutants for *capV*, *dncV*, or *cap2* has no effect on the plaquing efficiency against ICP2 or ICP3, leading to a phenotype similar to the WT strain (**Fig 2J**). This indicates that a functional CBASS system is necessary for toxigenic OueS to confer resistance against intestinal phages.

### OueS is associated with ToxR-mediated phenotypes

ToxR directly regulates the expression of *ompU*^61^. Consequently, we investigated the potential regulatory relationship between ToxR and OueS. Deletion of ToxR leads to an increase in biofilm production similar to a Δ*ompU* mutant (**Fig 3A**). Ectopic expression of either the *ompU*^C6706^ ORF or OueS^C6706^ in the Δ*toxR* strain restores biofilm repression to WT levels (**Fig 3A**). This OueS-mediated repression is specific as the WT phenotype could not be rescued in a Δ*toxR*::OmpU^mut^ strain or by ectopic expression of the OueS 5’-mut, 3’-mut or FL-mut alleles or the NS region within the *ompU* ORF (**Fig 3A**). Pairwise comparisons of WT and Δ*toxR*-pOueS^C6706^ relative to Δ*toxR* reveal that 42.69% of the ToxR regulated genes during biofilm formation overlap with the OueS transcriptome (155 of 363 genes; absolute FC ≥ 2.0, *q* ≤ 0.01; **Table S4**). The directionality of expression (activation Vs repression) in 149 of these 155 OueS-regulated genes (96%) is identical between the WT and Δ*toxR*-pOueS^C6706^ strains, strongly suggesting that OueS restores a significant portion of the regulatory changes observed with the loss of ToxR function in biofilms (**Table S4**). Overall, these findings indicate that the ToxR-mediated biofilm repression is due to downstream OueS expression and highlight a connection between the sRNA and ToxR regulome.

**Figure 3.**
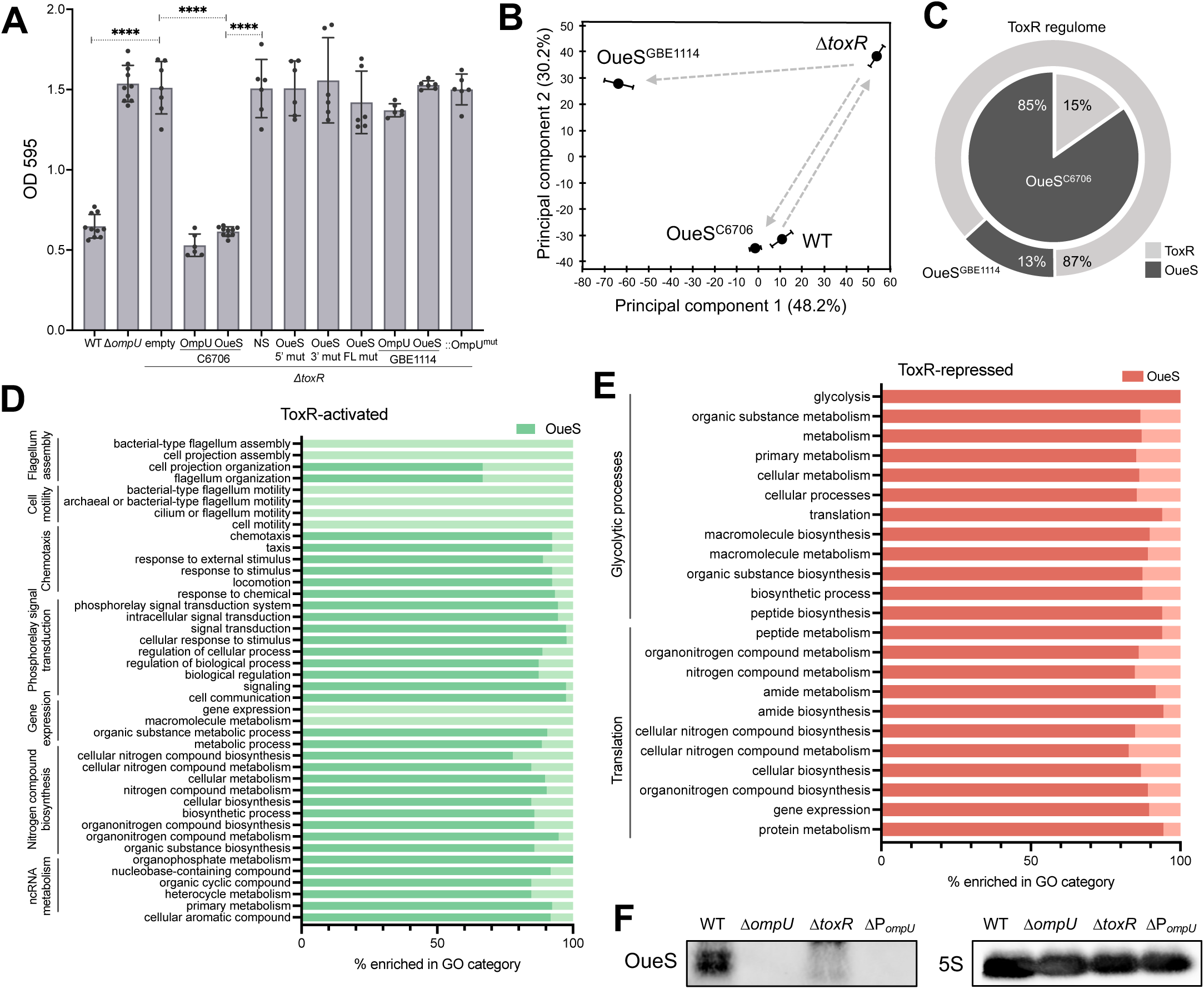
OueS expression is controlled by ToxR acting as its functional surrogate. (A) Ectopic expression of OueS represses biofilm formation in a Δ*toxR* mutant. Strains harbor either the empty vector or encode OmpU or OueS mutants and allelic variants. N ≥ 6. Two-tailed T-test *p*-value **** ≤ 0.0001. (B-E) OueS modulates the ToxR transcriptome under virulence-inducing conditions (AKI). (B) PCA plots show that a Δ*toxR* strain ectopically expressing OueS^C6706^ has a transcriptome resembling WT, unlike OueS^GBE1114^. (C) OueS^C6706^ (dark grey slice) modulates the expression of 84.7% (493 genes) of the ToxR virulence regulon (581 genes) (absolute FC ≥ 2.0; *q* ≤ 0.01), whereas OueS^GBE1114^ modulates 13.4% (dark outer circle). These do not include *ctxAB* or the *tcp* operon. (D) Functional enrichment analysis of the ToxR regulome under AKI conditions reveals that most of the functional categories that are (D) activated and (E) repressed by ToxR are primarily modulated by OueS (dark green and dark red, respectively). (F) Northern blot analysis reveals that OueS expression is controlled by ToxR via the *ompU* promoter. 5S rRNA probes are used as a loading control.

### Toxigenic OueS acts as a functional surrogate of ToxR

In order to further elucidate the role of OueS during the infectious process and its relationship with ToxR, we first examined their transcriptome under virulence-inducing conditions (AKI)^62,63^. Principal component analysis (PCA) of the RNA-Seq data shows a major shift in the transcriptome of the Δ*toxR* strain compared to WT by modulating the expression of 581 genes (400 activated, 181 repressed) (**Fig 3B; Table S5**). Ectopic expression of OueS^C6706^ in the Δ*toxR* mutant (Δ*toxR*-pOueS^C6706^) under AKI conditions results in a drastic shift in the transcriptome closely resembling the WT (**Fig 3B, Fig S4**). Interestingly, expression of the environmental allele OueS^GBE1114^ in the Δ*toxR* strain does not restore the transcriptome to WT levels, indicating that this overlap in the transcriptome is allele specific (**Fig 3B, Fig S4**). To examine the extent of OueS^C6706^ complementation, we determined the ratio of the fold change between the ToxR and OueS regulons [(*toxR*/WT) / (*toxR*-pOueS^C6706^/WT)]. Our analysis reveals that OueS modulates the expression of at least 493 of the 581 genes within the ToxR regulon (84.7%) under AKI conditions (**Fig 3C**; **Fig S4**; **Table S6**). Importantly, OueS does not modify expression of *ctxAB* or the TCP operon, which are activated by ToxR^18,22^ (**Fig 3C**; **Fig S4**; **Table S6**). We then performed enrichment analysis of the 581 genes of the ToxR regulome to determine the specific genes that OueS^C6706^ regulates based on functional categories. Our analyses show that OueS controls the ToxR-mediated upregulation of genes associated with chemotaxis, phosphorelay signaling as well as genes involved in nitrogen metabolism (**Fig 3D; Table S6**). On the other hand, OueS is responsible for the control of most of the ToxR-downregulated genes involved in glycolysis and translation (**Fig 3E; Table S6**). Finally, to determine whether the OueS-mediated modulation of the ToxR regulome is associated with the emergence of toxigenic clones of *V. cholerae*, we examined the transcriptome of Δ*toxR* strains ectopically expressing the OueS^GBE1114^ allele under AKI conditions. Pairwise comparisons [(*toxR*/WT) Vs (*toxR*-pOueS^GBE1114^/WT)] reveal that, unlike the toxigenic allele, OueS^GBE1114^ only modulates 13.4% (78 of 581 genes) of the ToxR regulome under AKI conditions (**Fig S4; Table S6**). Furthermore, OueS also modulates the OmpU transcriptome under virulence-inducing conditions (**Supplementary Text, Tables S7, S8**). Overall, the data shows that toxigenic OueS acts as a functional surrogate of ToxR under virulence-inducing conditions modulating the expression of 84.7% of its regulome.

### OueS is expressed from the *ompU* promoter

In order to identify a potential promoter of OueS, we used biofilm repression as a functional readout to initially monitor sRNA expression. First, we generated mutant strains where we cloned OueS together with varying lengths of its upstream region potentially harboring the promoter and integrated it at the native *ompU* locus in *V. cholerae* C6706. We did so in 10 bp increments (P_0_, P_10_, P_20_, … P_100_) from zero bp upstream of OueS (P_0_) to 100 bp upstream (P_100_) and monitored biofilm formation of these 10 strains. All the constructs repress biofilm formation similar to the WT, indicating that OueS does not have a native promoter within *ompU* (**Fig S5**). One likely scenario that explains this data is that OueS is co-transcribed with the *ompU* mRNA transcript from the *ompU* promoter. To verify this, we generated an *ompU* promoter mutant by deleting the 300 bp region immediately upstream of the *ompU* start codon, which encodes both the promoter and transcription start site^64^, generating the strain ΔP*_ompU_*, which retains an intact *ompU* and OueS coding region. Biofilm formation in ΔP*_ompU_* resembles the Δ*ompU* strain suggesting lack of OueS expression (**Fig S5)**. Furthermore, northern blot analyses reveal that, similar to Δ*ompU* or Δ*hfq* strains, OueS is undetectable in Δ*toxR* and ΔP*_ompU_* strains, demonstrating that OueS expression is controlled by ToxR via the *ompU* promoter (**Fig 3F**).

### OueS is required for successful intestinal colonization

OueS is essential for phenotypes associated with successful host colonization such as biofilm repression or resistance against intestinal phages and controls a large percentage of the ToxR regulon. To examine whether OueS plays a direct role in intestinal colonization, we performed competition assays using the infant mouse model of infection^65–67^. Inducer-dependent ectopic expression of the sRNA is not feasible in animal models. However, our data demonstrates that the strain expressing OueS directly from P*_ompU_* (P*_ompU_*-OueS^C6706^) exhibits an identical phenotype as the strain ectopically expressing the sRNA, making it suitable for *in vivo* intestinal colonization assays (**Fig S5**). Additionally, we generated isogenic strains encoding mutant OueS variants directly driven by P*_ompU_* at the native *ompU* locus, similar to the ectopic expression variants generated in **Fig 1H**. As above, we mutated every third nucleotide of the OueS sRNA locus that lies either exclusively within the *ompU* ORF (OueS 5’-mut), in the 3’ UTR (OueS 3’-mut) or across the full length of the sRNA (OueS FL-mut). The competitive fitness during intestinal colonization of these strains was determined against a *V. cholerae* C6706 *ΔlacZ* strain^31,66,67^. The ability of the various mutants to colonize the intestine relative to the WT strain was calculated by determining their competitive indices (CI). As expected, Δ*ompU* demonstrates a ∼1-log decrease in the CI of the small intestine (CI 0.06 – 0.14; **Fig 4A**). Interestingly, the *P_ompU_-* OueS^C6706^ strain rescues the colonization deficiency associated with the loss of OmpU and exhibits CI similar to WT (CI 1.05 – 1.29) (**Fig 4A**). These changes in colonization are specific to OueS as strains expressing the OueS 5’-mut (CI 0.15 – 0.44), 3’-mut (CI 0.17 – 0.48), FL-mut (CI 0.11 – 0.29) or OmpU^mut^ (CI 0.14 – 0.20) are not capable of restoring WT CIs (**Fig 4A**). Finally, the mutant with a deletion in the *ompU* promoter (ΔP*_ompU_*) shows a decrease in CI of the small intestine similar to the Δ*ompU* strain (CI 0.06 – 0.17; **Fig 4A**). Taken together, our results reveal that OueS is critical for successful intestinal colonization of toxigenic *V. cholerae*.

**Figure 4.**
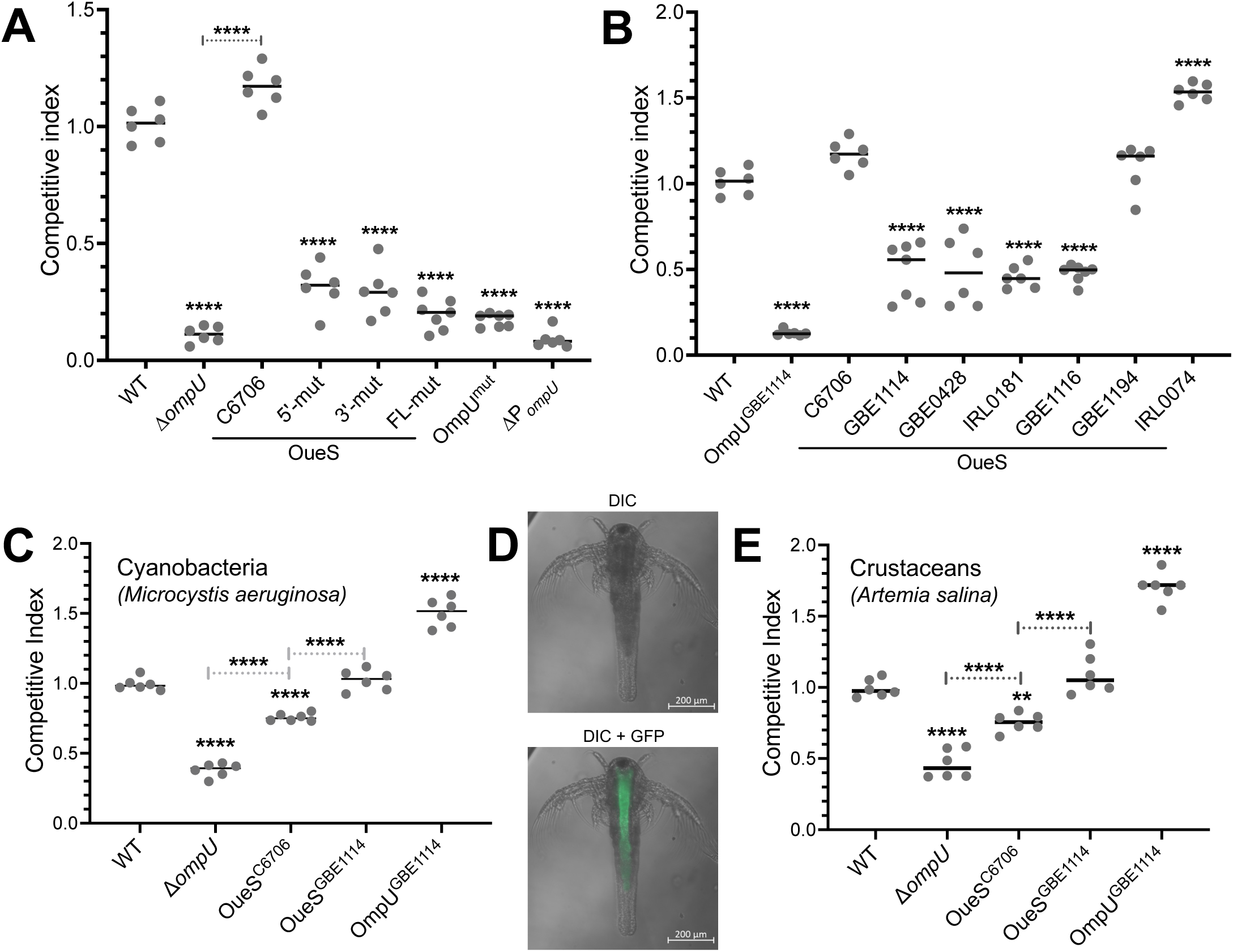
OueS leads to fitness trade-offs between intestinal colonization and environmental survival. *V. cholerae* strains were orogastrically inoculated in 3-5-day old CD-1 mice to perform competition assays (A, B). Data is represented as competitive indices (CI) between the test strain and a *V. cholerae* C6706 Δ*lacZ* control strain. (A) Expression of the toxigenic allele of OueS leads to successful intestinal colonization of *V. cholerae*. (B) OueS alleles from environmental strains expressed in the isogenic *V. cholerae* C6706 background leads to differential intestinal colonization indicating a role for OueS in the emergence of colonization potential. (C) Competition assays of *V. cholerae* strains in *Microcystis aeruginosa*. (D) Fluorescence microscopy image of *Artemia salina* infected with *V. cholerae* C6706 constitutively expressing GFP shows colonization of its gut space. (F) Competition assays of *V. cholerae* strains in *Artemia salina*. CIs in these reservoirs uncover the fitness trade-offs associated with the toxigenic OueS allele and the emergence of virulence traits in *V. cholerae*. n ≥ 6 per strain. *p*-values indicate significance compared to the WT strain, unless otherwise specified; one-way ANOVA * <0.01, ** < 0.001, *** < 0.0001, **** < 0.00001.

### Modular heterogeneity in OueS contributes to emergence of toxigenic strains

Our data demonstrates that specific allelic variations of OueS are essential for several virulence-associated phenotypes to emerge. Recently, we examined the natural genetic variability of *ompU* and identified 41 alleles of the gene from over 1600 sequences analyzed^41^. Sequence analysis of the OueS locus confirm the modular nature of the sRNA with extensive variation in the 5’ region encoded within the *ompU* ORF and its 3’ region consistently conserved (**Fig S6A**). We modeled the secondary structures of these alleles and identified a diverse set of conserved OueS structures, including the one from toxigenic strains (**Fig S6B**). To examine the roles of these alleles in host colonization, we constructed six isogenic mutant strains in *V. cholerae* C6706 encoding different OueS alleles covering its landscape of structural diversity, driven from P*_ompU_* at the native *ompU* locus (P*_ompU_*-OueS allele). Subsequently, we examined their competitive indices during intestinal colonization using the infant mouse model of infection. Our assays reveal a heterogenous colonization landscape mediated by the various OueS alleles. Mutants expressing the OueS allele from the environmental strain GBE1194 exhibit a CI similar to the WT strain (CI 0.8 – 1.2), whereas strains expressing OueS^IRL0074^ show higher CIs (CI 1.5 – 1.6; **Fig 4B**). On the other hand, isogenic mutants encoding OueS from environmental strains GBE1114, GBE0428, IRL0081, and GBE1116 exhibit a decrease in colonization that ranges between WT and Δ*ompU* (CI 0.45 – 0.56) (**Fig 4B**). Overall, our data indicates that **a)** the OueS allele encoded by toxigenic strains is a virulence preadaptation since some environmental strains (e.g., GBE1194) encode alleles that also lead to successful colonization, and **b)** OueS is important for the emergence of pathogenic potential in *V. cholerae*, as strains like GBE1114 encode alleles that lead to defective intestinal colonization.

Small RNAs modulate gene expression by directly binding to their target mRNAs ^68,69^. However, we did not find a direct correlation between the CI associated with a specific OueS allele and either its length (166-196 ntds) or its predicted secondary structures (**Fig S6**). Nonetheless, sequence variation analysis unequivocally reveals that OueS is bimodular, comprising of variable *ompU*-encoded 5’ modules and a conserved 3’ one, the former directly contributing to the emergence of pathogenic *V. cholerae* from environmental populations. To correlate OueS modules with their transcriptional profiles, we used the infant mouse intestinal colonization data to classify the OueS alleles into toxigenic-like (GBE1194, IRL0074) and environmental-like (GBE1114, GBE0428, GBE1116, IRL0181) variants (**Fig 4B**). Analysis of the AKI transcriptomes of representative toxigenic-like (C6706, GBE1194) and environmental-like (GBE1114, GBE1116) OueS allele-expressing strains identified 96 genes (57 repressed and 39 activated) exclusively regulated by the toxigenic-like OueS alleles (**Fig S7; Table S9**). However, we did not find significant functional enrichment among the 96 genes indicating that no single group of genes or pathway is over-represented in the dataset. This suggests a complex scenario with potential regulatory redundancies and compensatory networks associated with the emergence of these phenotypes.

### Allelic variability of OueS leads to fitness trade-offs between human infection and environmental survival

*V. cholerae* is a natural inhabitant of aquatic biomes and can be found associated with reservoirs that critically affect disease transmission^70^. To examine the role of OueS in *V. cholerae* environmental survival and the potential fitness trade-offs associated with its toxigenic allele, we determined the CIs of OueS mutants during the colonization of two model environmental reservoirs established in the lab: the cyanobacteria *Microcystis aeruginosa* and the crustacean *Artemia salina* (**Fig 4C, D and E**)^70^. We performed competition assays in both model systems following a similar approach as above, where we compared the colonization potential of test strains (*lacZ*^+^) against a *V. cholerae* C6706 Δ*lacZ* control strain. Our results indicate that OmpU is important for colonization of both *M. aeruginosa* (**Fig 4C**) and *A. salina* (**Fig 4E**) as a Δ*ompU* mutant exhibits a ∼6-fold reduction in its CI (0.39 and 0.43, respectively). Expression of toxigenic OueS (OueS^C6706^) leads to a significant increase in CI in both the cyanobacteria (CI 0.75) and crustacean (CI 0.76) model systems when compared to Δ*ompU* (**Fig 4C, E**). Interestingly, the CI of strains expressing an environmental OueS allele (OueS^GBE1114^) closely resembles that of the WT strain (CI 1.05 – 1.06) in both reservoirs. Furthermore, mutant strains expressing the environmental OmpU allele (OmpU^GBE1114^), which encodes both the environmental porin and sRNA, outcompete the WT strain encoding its toxigenic versions in both reservoirs (CI 1.51 - 1.67). Overall, our results indicate that OmpU is important for colonization of environmental reservoirs in a OueS-dependent manner and its toxigenic allele leads to fitness trade-offs when compared to its environmental counterpart.

## DISCUSSION

Emerging and re-emerging infectious diseases pose immense health and economic risks^71^. To date, the evolutionary drivers that facilitate certain strains within a population to develop virulence traits and emerge as human pathogens remain largely unknown. This critically hinders our ability to develop effective strategies (e.g., policies, therapeutics, vaccines, etc.) against facultative pathogens in a timely fashion. Previously, we proposed that the presence of allelic variations in core genes might explain the emergence of virulence traits in toxigenic *V. cholerae* isolates^31^. The gene coding for the major outer membrane protein OmpU encodes these allelic variations in the form of virulence adaptive polymorphisms (VAPs) and plays an integral role in pathogen emergence^31,41^. In this study, we demonstrate that the *ompU* ORF encodes a sRNA with a unique modular structure that plays a central role in the emergence of virulence traits in *V. cholerae* (**Fig 5A**). The 5’ end of the OueS locus is highly diverse and is encoded within the *ompU* ORF, whereas its 3’ terminus is conserved and lies in the UTR of the ORF. This atypical location confers a bimodular structure to OueS that generates allelic variants contributing to the emergence of virulence-associated phenotypes in some strains, a property that is unique among known sRNAs. Investigating the prevalence of this configuration will shed light on the role of sRNAs in the evolution of bacterial pathogens (46-48, 62). The expression of OueS is driven from P*_ompU_* and is regulated by ToxR, potentially generating one single transcript (**Fig 5A**). It remains to be determined how the combined *ompU*-OueS transcript is processed to generate both the porin and sRNA, which could be mediated by ribonucleases such as RNase E or others^48,72^. Furthermore, OueS expression peaks during the early stationary phase, drastically decreasing at later growth stages by yet unknown mechanisms (**Fig 1G**). Findings that elucidate whether the expression and processing rates of OueS are modulated in response to specific host signals during the infection process will provide critical insights into the mechanisms governing *V. cholerae* spatiotemporal colonization dynamics.

**Figure 5.**
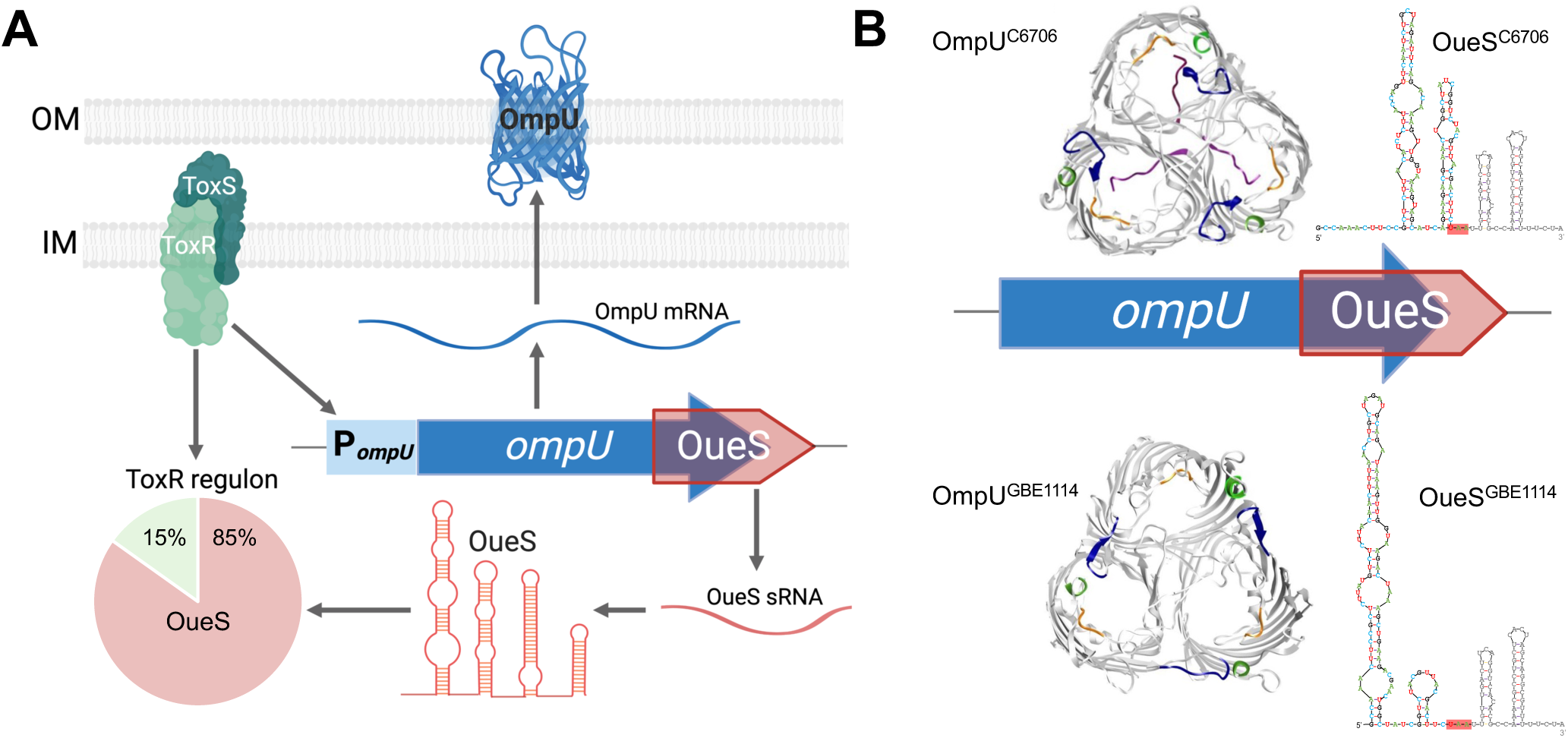
The *ompU* ORF encodes a modular porin and sRNA. (A) The virulence regulator ToxR controls the expression of hundreds of genes including the *ompU* ORF via P*_ompU_*. The ORF encodes both **a)** the OmpU porin (blue) and **b)** the OueS small RNA (red), which encompasses its 3’ terminus and UTR. The toxigenic allele of OueS acts as a functional surrogate of ToxR by modulating the expression of ∼85% of its regulome. (B) *V. cholerae* encodes numerous alleles of *ompU* with distinct modular versions of OmpU and OueS structurally and functionally. Modularity in the porin and the sRNA enables the emergence of phenotypes essential for host colonization such as antimicrobial resistance, stringent biofilm regulation, and host transmission dynamics.

We determined that the OueS allele from toxigenic strains controls phenotypes essential for host colonization and survival. For instance, it inhibits biofilm formation by repressing RyhB, a well-characterized sRNA controlled by Fur (**Fig 2C-E**)^51,53^. Interactions between sRNAs to modulate gene expression is being increasingly recognized as a critical component governing cell morphology and behavior^73–77^. For instance, expression of the major stress response regulator *rpoS* in *Escherichia coli* is reciprocally modulated by interactions between two sRNAs, ArcZ and CyaR^76^. Similarly, interactions between sRNAs modulate cell division and ethanol tolerance in some non-pathogens^73,74^. To our knowledge, our finding that OueS regulates RyhB in an allele-dependent manner is the first identified example of a sRNA-sRNA interaction that directly affect host colonization and the emergence of pathogenic traits in bacteria. Recent high throughput studies demonstrate potential to identify interactions between sRNAs on a whole genome scale such as RNA sponges^75,77^, opening avenues to unravel fascinating aspects of the intricate regulatory network in bacteria.

Bacteriophages are a key driver in the emergence of pandemic *V. cholerae* and play a critical role in its intestinal survival and colonization, environmental prevalence, and seasonality of cholera epidemics^30,55–57,78,79^. Our data demonstrates that expression of the toxigenic allele of OueS confers resistance to two phages intimately associated with cholera dynamics: ICP2 and ICP3, both of which were isolated from stool samples of cholera patients (**Fig. 2I**)^80^. We determined that OueS mediates phage resistance via activation of the CBASS abortive infection system. The CBASS system was previously shown to confer resistance to circulating variants of the ICP3 phage and thus, our findings provide a molecular and evolutionary mechanism explaining this phenotype^81^. Interestingly, the OmpU porin, which is encoded in the *ompU* ORF together with OueS, is the cell receptor for ICP2^80^. Our data reveals an ingenious mechanism developed by the cell to defend against phage infections by co-transcribing a sRNA that confers phage resistance along with the porin that functions as the receptor. Given how commonly phages utilize porins as cell receptors, we contend that examining the ubiquitousness of this arrangement will unveil fascinating nuances and novel mechanisms used by bacteria to resist viral infections. A cocktail of ICP phages have been proposed as a therapeutic intervention to prevent host colonization by the bacterium^78^. Therefore, elucidating the associated resistance mechanisms will shed new light in the arms race between bacteria, phages and their receptors, which have direct implications in preventing infections and cholera outbreaks.

The transcriptional regulator ToxR plays a key role in virulence and other physiological processes by modulating the expression of hundreds of genes in *V. cholerae*^20,21^. Surprisingly, ToxR only binds directly to the promoter of 39 genes^21^. Here, we provide a mechanism that elucidates this conundrum as 84% of ToxR regulome can be explained by the expression of toxigenic OueS via activation of the OmpU promoter by ToxR (**Fig 3**, **Fig S4**). It is possible that this is not an isolated example and other sRNAs act as functional surrogates of virulence regulators, suggesting a unique mechanism among bacterial pathogens. Elucidating this will shed light on the regulatory cascades of other DNA-binding regulators and provide novel downstream targets for therapeutic interventions. Interestingly, OueS does not modulate expression of CT or TCP, indicating a compartmentalized role of the sRNA within the ToxR regulome (**Fig 3**). Furthermore, the transcriptome of the environmental OueS^GBE1114^ allele does not overlap with the ToxR regulome which highlights the close relationship between the evolution of the sRNA and the emergence of virulence traits in *V. cholerae*. This is further evidenced by the role that allelic variability in OueS plays in intestinal colonization, where some alleles like the toxigenic one confer critical competitive advantages during infection (**Fig 4**).

*V. cholerae* is a natural inhabitant of aquatic biomes and is often associated with reservoirs that contribute to their environmental survival^70,82^. We uncovered a fitness trade-off associated with the expression of OueS alleles and colonization of environmental reservoirs. Our results show that the toxigenic allele of OueS leads to fitness costs during environmental survival as demonstrated by the strain encoding the environmental allele exhibiting a higher CI during colonization of both *A. salina* and *M. aeruginosa* (**Fig 4**). It is tempting to hypothesize that these selective pressures function as bottlenecks restricting the emergence of pathogenic traits in environmental strains of *V. cholerae*. These fitness costs might explain the limited distribution of toxigenic isolates of *V. cholerae* and other human pathogens. Future research that delineates how the various biotic and abiotic factors in the estuarine environment influence *V. cholerae* colonization dynamics will be critical to understand the evolution of its virulence traits and host transmission potential.

Modularity has become a central concept in biology, stemming from the idea that biological systems are intricately linked networks that are typically divided into sets of interacting modular subsystems^83^. Modularity spans many scales in biological systems and provides a cohesive mechanistic framework to explain biological complexity^84–87^. Recently, we determined that the porin OmpU contains modular protein domains that uniquely contribute to the emergence of antimicrobial resistance in certain alleles^41^. Here, we unveil a new layer in the complex modular nature of the *ompU* ORF and demonstrate that OueS also displays modular characteristics (**Fig 5B**). OueS is a bimodular sRNA comprising of **a)** a variable 5’ module encoded within the *ompU* gene that contributes to diversity and emergence potential, and **b)** a highly conserved 3’ module downstream from the *ompU* ORF that likely plays a role in sRNA stability and target recognition by binding Hfq. Taken together, our results indicate that the *ompU* ORF encodes both a porin and a sRNA, and their modular nature enables the emergence of phenotypes essential for host colonization such as antimicrobial and phage resistance, stringent biofilm regulation, and host transmission dynamics (**Fig 5**). We propose a scenario where a limited network of highly variable modular genes like *ompU* could lead to phenotypic heterogeneity in a population including the emergence of virulence-associated traits. Recently, we analyzed over 1,800 *V. cholerae* genomes and identified several genes that possess similar patterns of allelic diversity^40^. We found that the allelic variations associated with toxigenic strains confer a non-linear competitive advantage during intestinal colonization acting as a critical bottleneck that helps decipher the isolated emergence of toxigenic *V. cholerae*^40^. Elucidating the molecular processes and interaction dynamics between these genes will yield critical insights into the evolution of facultative pathogens. Overall, our findings provide mechanistic insights into the emergence of pathogenic traits in bacteria unraveling the hidden genetics and fitness costs associated with pathogen emergence.

## METHODS

### Bacterial strains, media, and growth conditions

All *V. cholerae* strains used in this study are derivatives of the seventh pandemic El Tor biotype strain C6706, first isolated from a patient in Peru in 1961^90^. *E. coli* S17 λ-*pir* strains harboring a chromosomally integrated RP4 plasmid and *tra* genes^91^ were used for conjugation of the deletion constructs into *V. cholerae* as well as for routine cloning purposes. Phage propagation was performed on *V. cholerae* AC6169, a BSL1 derivative of the El Tor strain E7946. The following media were used for bacterial cultivation: Luria Bertani (LB) broth (10 g l^-1^ tryptone, 5 g l^-1^ yeast extract, 5 g l^-1^ NaCl); Biofilm assays were performed in tryptone broth (tryptone 10 g l^-1^, NaCl 5 g l^-1^); AKI broth was used for virulence-inducing conditions^62,63^ (15 g l^-1^ Bacto^TM^ peptone, 4 g l^-1^ yeast extract, 5 g l^-1^ NaCl, 0.03% w/v sodium bicarbonate). Plates were used at a final concentration of 1.5% agar. Top agar for phage assays contained LB broth + 0.7% agar. Where necessary, the following antibiotics or supplements were used: streptomycin 1000 µg ml^-1^, kanamycin 45 µg ml^-1^, polymyxin B 50 U ml^-1^, carbenicillin 50 µg ml^-1^, 2,2’-bipyridyl 100 µg ml^-1^, FeCl_3_ 100 µg ml^-1^, 5-bromo-4-chloro-3-indolyl-ß-D-galactopyranoside (X-gal) 40 µg ml^-1^, and arabinose (0.01%). Reagents were procured from Sigma.

For routine culture, *E. coli* and *V. cholerae* were grown in LB broth at 37 °C with aeration (200 rpm). Biofilm and growth curve assays were performed in 96-well flat bottom, tissue culture-treated plates (CytoOne) incubated at 37 °C.

### Cloning, vectors, and strain construction

In-frame deletion mutants of *ompU*, *toxR, ryhB*, and *lacZ* were constructed via homologous recombination^92^. Briefly, approximately 500 bp fragments flanking the gene of interest to be deleted were PCR amplified, cloned as *Bgl*II-*Not*I (upstream fragment) or *Not*I-*Eco*RI (downstream fragment) in tandem into the *V. cholerae* suicide vector pKAS154 ^92^, and electroporated into *E. coli* S17λ-*pir*^91^. Deletion constructs of genes of the CBASS system (*capV*, *dncV*, *cap2*) were generated using the HiFi DNA Assembly (NEB) to prevent polar effects of the deletions within the operon. *E. coli* strains harboring the deletion constructs were then conjugated with *V. cholerae* C6706, and allelic exchange was carried out with appropriate antibiotic selection, as described^92^. Positive clones were first screened by colony PCR and then confirmed by sequencing the deletion loci from genomic DNA using flanking primers. The *ompU* promoter mutant ΔP*_ompU_,* was generated in a similar manner by amplifying the regions immediately flanking the 300 bp locus right before the *ompU* start codon, generating a deletion construct in pKAS154, followed by allelic exchange into C6706, as described above. The OmpU^mut^ strain with an intact OmpU amino acid sequence but a mutated OueS and driven by P*_ompU_* was generated by making wobble mutations across the entire length of the OueS sequence (**Fig S2**). The fragment was then synthesized commercially (Twist Biosciences) along with 500 bp of the *ompU* downstream region and cloned as a *Bgl*II-*Eco*RI fragment into pKAS154. Allelic exchange with WT C6706 was then performed as described above.

Ectopic OueS-expressing clones and non-specific *ompU* locus clones were generated by PCR amplification from the appropriate genomic DNA template using specific primers listed in **Table S10**. The amplicons were then cloned as *Eco*RI-*Hin*dIII fragments into pBAD22. The OueS modular mutants were synthesized commercially (Twist Biosciences) and then cloned into pBAD22 as above. After introduction into *E. coli* by electroporation and sequence confirmation of the inserts, the plasmids were electroporated into the appropriate *V. cholerae* strains, followed by selection on carbenicillin-containing LB plates. For strains harboring the empty vector, plasmid pBAD22 was electroporated into the appropriate *V. cholerae* strain and selected as above. Ectopic *ompU*-expressing clones were generated in a similar manner.

Chromosomal promoter fusions were constructed as described above for the deletion constructs. The ∼500 bp regions upstream of the *ompU* ORF including P*_ompU_ was* first cloned as a *Xba*I-*Sac*I fragment into pKAS154 to generate plasmid pDBS467. Subsequently, the various OueS constructs: OueS 5’-mut, 3’-mut, FL-mut fragments (commercially synthesized; Twist Biosciences), and the various OueS alleles (amplified from the respective genomic DNA) were cloned into pDBS467 downstream of P*_ompU_* as a *Sac*I-*Eco*RI fragment. Allelic exchange was then performed as described above.

All deletion and complementation constructs in *E. coli* were Sanger sequenced before introduction into *V. cholerae*. Genomic DNA was isolated from all *V. cholerae* deletion and chromosomal fusion strains, and the region(s) of interest amplified using flaking primers for confirmation by Sanger sequencing.

### RNA isolation and transcriptome studies

For Northern blots and RACE studies, RNA was isolated using the hot-phenol extraction method at the appropriate growth stage and conditions^93–95^. Briefly, the cell pellets were harvested and resuspended in cell lysis solution (0.5% SDS, 20 mM sodium acetate, 10 mM EDTA), followed immediately by the addition of 500 µL of acid-phenol:chloroform (Ambion) preheated to 65 °C and vortexed. After incubation at 65 °C for 10 minutes, the samples were centrifuged at 21,000 x g for 10 minutes and the aqueous layer reextracted twice with acid-phenol:chloroform. The aqueous phase was then extracted twice with chloroform and precipitated overnight at −80 °C with 2.5 volumes of ice-cold absolute ethanol. The total RNA was then pelleted and washed with cold 70% ethanol, pellets dried and resuspended in 100 µl RNase-free water containing 40 U murine RNase inhibitor (New England Biolabs). Following quantification by Nanodrop, 50 µg of the total RNA was DNase-treated (Ambion RNase-free DNase) following manufacturer protocols. The RNA was reextracted as above and precipitated with 1/10^th^ volume 3M sodium acetate and cold absolute ethanol followed by a cold 70% ethanol wash. Final resuspension of the dried RNA pellets was in 50 µL of RNase-free water containing RNase inhibitor (as above). Size fractionation for RACE studies was performed by running 10 µg of hot phenol-isolated RNA on a precast 1.25% MOPS agarose gel (Lonza) in 1x MOPS buffer (Quality Biological) with the Riboruler RNA ladders (Thermo) for size determination. Gel extraction was performed using the Zymoclean gel RNA recovery kit (Zymoresearch).

For transcriptome studies, RNA was isolated using a commercial kit (DirectZol RNA Miniprep kit, Zymo Research) following manufacturer instructions. DNase treatment, rRNA depletion, library preparation and Illumina NextSeq2000 sequencing (2×51 bp reads) was performed at SeqCenter, LLC (Pittsburgh, PA). Demultiplexing, quality control and adapter trimming was performed with bcl-convert (v3.9.3). CLC Genomics Workbench (Qiagen) was used for subsequent data processing. Briefly, paired Illumina reads were mapped to the El Tor reference genome (GenBank AE003852.1-Chromosome I; AE003853.1-chromosome II) using the following settings to generate the transcripts per million bases (TPM) values: mismatch cost 2, insertion / deletion cost 3, length / similarity fraction 0.8, global strand specific alignment. Subsequent pairwise analysis between two datasets of interest (absolute fold change ≥ 2.0 filtered on average expression for FDR correction ≤ 0.01) were used for downstream analysis. Functional enrichment analysis of the transcriptome data was performed by the Panther Overrepresentation Test against the Biological processes dataset of the GO Ontology Database (Released 07-01-2022)^88,89^.

### Biofilm assays

The ability of the strains to form biofilms on surfaces was determined using the crystal violet-based assays in 96-well polystyrene plates, optimized from published protocols^96^. Briefly, the strains from LB plates were grown in tryptone broth with carbenicillin for 14-16 hours at 37 °C with aeration. The cultures were then diluted 1:500 in fresh tryptone broth supplemented with carbenicillin and arabinose and 200 µL dispensed in wells of the 96-well plates. After static incubation for 24 hours at 37 °C, the cultures were poured off and plates gently washed three times with water to remove unattached cells. The surface attached cells were then stained with 225 µL per well of crystal violet (0.1% in water) for 20 minutes at room temperature and washed thoroughly with water. The attached stain was subsequently eluted with 225 µL per well of 50% acetic acid for 15 minutes and quantified by reading the absorbance in a Tecan Sunrise plate reader at 595 nm. To examine the effect of iron on biofilm formation, TB media was supplemented with either 100 µM FeCl_3_ (MP Biomedicals) for iron repletion or 100 µM 2, 2’-bipyridyl (Sigma) to deplete residual iron. Growth and processing of biofilms was performed as described above. For transcriptome studies, 24-hour biofilms were harvested by scrapping off attached cells into fresh media and used for total RNA isolation.

### Northern blot assays

For northern blot assays, total RNA isolated using the hot-phenol method (see above) were used. Briefly, 6 - 10 µg RNA per lane was first resolved on a 1.2% agarose gel (Lonza) in 1 x MOPS buffer (Quality Biological) and transferred to BrightStar Plus positively charged nylon membranes (Invitrogen) by electroblotting (Trans-Blot SD transfer cell, BioRad). Following transfer, the membranes were UV crosslinked (UV Crosslinker, Fisherbrand), prehybridized for 5 hours (ULTRAhyb Ultrasensitive Hybridization buffer, Invitrogen) in roller bottles in a UVP Hybridizer Oven (Analytik Jena) at 42 °C. Commercially synthesized 5’ biotinylated probes (10 pmoles of 40-mer probes; Eurofins Genomics) for the RNA targets were used for hybridization for 15 hours at 42 °C, washed with low- and high-stringency buffers (Invitrogen), and developed using the Chemiluminescence Nucleic Acid Detection kit (Thermo), following manufacturer instructions. Developed blots were imaged on a Bio-Rad ChemiDoc MP imaging system.

### Rapid Amplification of cDNA Ends (RACE)

The 5’ and 3’ termini of the sRNA OueS were determined by the RACE assay (SMARTer RACE 5’/3’ kit, Takara Bio USA), adapted for non-poly-A-tailed RNA. Kit components were used, unless otherwise mentioned. Briefly, total RNA was separated on a 1.2% MOPS agarose gel and the 100-500 ntd region was gel extracted (Zymoclean Gel RNA recovery kit). This size fractionated RNA was used as the template for both 5’ and 3’ RACE studies.

For 5’ RACE, first strand cDNA synthesis was performed by random priming followed by annealing the SMARTer IIA oligonucleotide to the 5’ ends of the newly synthesized cDNA fragments. This was followed by PCR amplification using a forward primer specific to the 5’-attached oligonucleotide and a reverse primer within the *ompU* gene (DB196: 5’ GATTACGCCAAGCTTtcgtaacgtagaccgatagccagttcgtct 3’). The fragments were then cloned non-directionally using the A-overhangs generated during the final PCR step and electroporated into *E. coli* Stellar Competent cells (Clontech) and selected on carbenicillin plates. Plasmids from positive clones were then sequenced to identify the 5’ terminus of OueS. For 3’ RACE, the size-fractionated total RNA was poly-A-tailed using the *E. coli* Poly(A) polymerase following manufacturer instructions (New England Biolabs). cDNA was then synthesized using a primer containing a complementary poly-T sequence and a 5’ overhang (DB164: 5’ GGGGGGGGAATTCTTTTTTTTTTTT 3’). The newly synthesized cDNA was used as template for a second strand synthesis using an *ompU*-specific primer that binds inside the 3’ end of the *ompU* ORF (DB195: 5’ GATTACGCCAAGCTTccgctcttacatctcttaccagttcaatctgctag 3’). The amplicons were then gel purified, and PCR amplified using the 3’ *ompU*-specific primer and a primer targeting the 5’ overhang of the poly-T primer (DB164) used in the first step of the process. The resulting amplicons were then cloned non-directionally into vectors with a T-overhang and electroporated into *E. coli*. Subsequently, plasmids from positive clones were sequenced to define the 3’ terminus of OueS.

### Prediction of sRNA secondary structure and target interaction

The secondary structures of the WT and mutant versions and alleles of OueS were determined using mFold on the UNAFold web server (RNA Folding Form V2.3)^97^ using default parameters. Structures with the least predicted ΔG values were selected. Interactions between OueS and its target RNAs such as RyhB were determined using IntaRNA (Freiburg RNA Tools)^98^ using default parameters.

### Growth kinetics under iron limitation

Overnight LB-grown cultures were diluted 1:1000 in LB broth with or without supplementation with FeCl_3_ (iron-replete) or 2,2’-bipyridyl (iron-deplete). 200 µL aliquots of the cultures dispensed in flat-bottom 96-well plates (tissue culture-treated, CytoOne) were then monitored for OD_595_ every 30 minutes for 24 hours at 37 °C in a Tecan Sunrise plate reader. Each biological replicate had at least six technical replicates and the data for each condition were averaged across technical and biological replicates corrected to the baseline (LB) absorbance values.

### Bacteriophage assays

Phages ICP2 and ICP3 were propagated on *V. cholerae* AC6169, a modified non-toxigenic El Tor strain that has a phase-locked expression of the O1 antigen (A. Camilli). Briefly, log phase cells (OD600 of 0.5-0.7) in LB broth were infected at a multiplicity of infection of 0.1 with the respective phages and allowed propagation for 3-5 hours, 200 RPM at 37 °C. Phage lysates were harvested by centrifugation and filtered through a 0.2 µ filter before use. Top agar overlays were performed using media supplemented with carbenicillin and arabinose of the different strains and the phage dilutions were spot titrated. The efficiency of plating (EOP) was calculated for ICP2 and ICP3 lysates: EOP = (titer on test strain / titer on WT C6706).

### Infant mouse competition assays

All animal procedures were conducted in accordance with the guidelines and protocols approved by the Institutional Animal Care and Use Committee (IACUC) at the University of Central Florida (#IPROTO202300049). For *in vivo* animal experiments, three-to-five-day-old CD-1 ® IGS mice (Strain code 022; Charles River Laboratories) of both genders were randomly selected from mixed litters for all animal experiments. Infant mice were housed with the mothers, monitored under the care of full-time staff, and weaned 2 hours prior to infection.

Intestinal colonization competition assays in three-to-five day-old infant mice were performed with LB-grown *lacZ*^+^ cells of the WT, deletion mutants and OueS alleles competed against an isogenic WT strain harboring a *lacZ* deletion, essentially as described^31,66,67^. The colonization efficiencies of the different strains in the small intestine harvested at ∼22 hours post infection are represented as competitive indices^66^. Briefly, strains were inoculated from fresh LB plates into LB broth, incubated for 12 hours at 37 °C with aeration, and diluted 1:1000 fold in fresh LB broth. The test (*lacZ*^+^) and control (*lacZ*^-^) strains were mixed 1:1 to obtain ∼10^6^ CFU/mL. Fifty microliters of the bacterial mixture, corresponding to ∼10^5^ CFUs, were then used to intragastrically inoculate 3-5-day-old infant mice and maintained at 30 °C for 22 hours. The exact input CFU numbers were determined by plating dilutions on LB-Sm-X-Gal plates. Approximately 22 hours post infection, the mice were euthanized, and their small intestines transferred into 4 mL LB-10% glycerol stored on ice. The samples were then processed for CFU by homogenizing the tissues, serially diluting in 1x PBS (pH 7.4) and plating appropriate dilutions on LB-Sm-X-Gal plates. *In vitro* competition experiments were performed in parallel by inoculating 50 µL of the input CFU in LB-Sm broth and incubating for 22 hours at 37 °C with aeration. Samples were then serially diluted and plated as for the *in vivo* samples above. Results are presented as the competitive index (CI), which is the ratio of the intestinal *lacZ*^+^ CFU (blue colonies) to *lacZ*^-^ CFU (white colonies) normalized to the input CFU ratio and *in vitro* competition CFU ratio.

### Colonization of environmental reservoirs

Bacterial inoculum for infections of the environmental hosts were prepared essentially as described for the mice colonization assays but without the Evan’s blue dye. Infections for fluorescent visualization were performed with a GFP-tagged *V. cholerae* C6706 strain and images obtained using a Carl Zeiss Axio Observer 7 Inverted Microscope. **a) *Artemia salina***. Briefly, eggs of *Artemia salina* were hatched in artificial sea water supplemented with 2% sodium chloride (ASW), incubated at room temperature and aeration for 48 hours. Infections were performed in 96-well flat bottom plates with each well containing 200 µL ASW with 20-30 individual crustaceans and 10^7^ CFU/mL of bacterial mixture. After 48 hours incubation at room temperature, samples were washed twice in ASW and either homogenized for determining CIs (as above) or used directly for microscopy. **b) *Microcystis aeruginosa***. We followed modifications of previously established protocols^99^: *Microcystis aeruginosa* (UTEX LB 2385) were incubated in flasks containing BG-11 media (Gibco) under a 12 h light-dark cycle with aeration (100 rpm) for about 14 days. The cyanobacterial cells then were centrifuged, resuspended in media and co-cultured at a final OD_700_ of 0.6 with 10^5^ CFU/mL *V. cholerae* cells in a 200 µL reaction volume. CIs were determined after 24 hours incubation by plating dilutions on LB-Sm-X-Gal plates, as above.

### Statistical analyses

All data were analyzed for statistical significance using Prism 10 software (GraphPad Prism), except for the transcriptome data that was analyzed within the CLC Genomics Workbench (Qiagen). Unless otherwise indicated, all experiments were performed with at least five independent biological replicates (n ≥ 5) and the aggregate data (mean or median ± s.d.) are shown.

## DATA AVAILABILITY

RNA-Seq data have been deposited in the GEO database: accession numbers GSE272767 (GSM8410380 – GSM8410418) and GSE272640 (GSM8408188 - GSM8408195).

## ACKNOWLEDGEMENTS

The authors would like to thank Dr. Andrew Camilli (Tufts University School of Medicine) for generously providing the ICP phages and *V. cholerae* AC6163 strain. This work was supported by a National Science Foundation (NSF) CAREER award (#2045671) and a Burroughs Wellcome Investigator in the Pathogenesis of Infectious Disease (#1021977) to SAM.

## AUTHOR CONTRIBUTIONS

SAM designed research. DB and CC performed research. DB, CC, and SAM analyzed data and wrote the manuscript.

## SUPPLEMENTARY MATERIAL

### Supplementary Text

#### OueS regulates the OmpU transcriptome under virulence inducing conditions

OmpU is highly expressed during the infection process and is critical for intestinal colonization^7,19,26^. We examined the role of OmpU and OueS in the context of virulence-associated conditions. To accomplish this, we first analyzed the transcriptomes of the WT, Δ*ompU* and Δ*ompU*-pOmpU^C6706^ strains under virulence-inducing (AKI) conditions^62^. Our analyses reveal that OmpU regulates a much larger proportion of genes under AKI conditions than during biofilm formation, modulating the expression of 947 genes (560 activated and 379 repressed), compared to 270 genes (181 repressed, 89 activated) in biofilms (**Table S7**). Ectopic expression of OmpU in Δ*ompU* decreases the number of differentially expressed genes to 170 from the 947 genes in Δ*ompU* (82.04% recovery of the Δ*ompU* transcriptome to WT levels) (**Table S7**). Furthermore, most of the OmpU-modulated genes under AKI conditions are RpoE-independent as only 158 of the 947 (16.68%) are shared with the RpoE regulon under AKI conditions, suggesting a much lesser role of the envelope stress response during the infection process (**Table S7**). Additionally, OueS modulates the expression of 455 of the 947 genes (48%) of the OmpU transcriptome under AKI conditions suggesting an integral role of OueS in this process (**Table S8**).

### Supplementary Figures

**Figure S1. Genes regulated by OmpU during growth in biofilms.** (A) Transcriptomes of biofilm-derived cells were visualized by principal component analysis. Ectopic expression of OmpU and OueS result in a similar shift in the datasets compared to the WT and Δ*ompU* strains. Functional enrichment of the OmpU-regulated genes was performed by the Panther Overrepresentation test (GO Biological process)^88,89^ to classify the OmpU-repressed (B) and activated (C) datasets. OmpU primarily represses iron-related genes and activates those involved in viral defense.

**Figure S2. OueS mutants and alleles**. (A) Sequence conservation of the *ompU* ORF encompassing the OueS 5’ locus before (WT) and after codon wobble mutations (OueS 5’-mut, 3’-mut and FL-mut) are shown to demonstrate an unaltered protein coding sequence. The 166 ntd OueS sequence from (B) C6706 (1′G −40.0), (C) OueS 5’-mut (1′G −40.7), (D) OueS 3’-mut (1′G −24.6), and (E) OueS full length-mut (1′G −26.5) were used to generate their predicted secondary structures on mFold (unafold.org). The structures requiring the least free energy (ΔG) are shown. Mutations significantly alter the predicted structures of the OueS mutants (C, D, E), compared to the WT OueS^C6706^ (B). (F) Pairwise alignment of the clinical OueS^C6706^ and environmental OueS^GBE1114^ alleles highlight variations arising from sequence differences in the sRNA region within the *ompU* ORF, resulting in a bimodular sRNA.

**Figure S3. Altered iron concentrations do not affect planktonic growth**. The planktonic growth patterns of the WT, Δ*ompU* cells ectopically expressing either the WT or mutated versions of OueS in (B) tryptone broth (baseline iron levels) and under (C) iron-replete (100 µM FeCl_3_) or (D) iron-depleted (100 µM 2,2’-bipyridyl) conditions. Growth patterns are similar between the strains in each condition irrespective of iron concentration. Biofilm formation of strains ectopically expressing the WT or mutant versions of OueS in the Δ*ompU* background were examined under iron-replete (100 µM FeCl_2_; dark grey bars) or iron-depleted (100 µM 2,2’-bipyridyl) conditions, compared to the basal iron conditions (TB media; light grey bars).

**Figure S4. OueS^C6706^ complements loss of ToxR under virulence-inducing conditions**. Pairwise comparisons of AKI transcriptomes of the WT strain and the *toxR* mutants expressing alleles of OueS reveals that (A) OueS^C6706^ restores the Δ*toxR* transcriptome to closely resemble the WT, whereas (B) OueS^GBE1114^ does not. All genes in this dataset have a *q*-value of ≤ 0.01 and are listed in Supplementary Table 6.

**Figure S5**. **OueS is expressed from the *ompU* promoter.** Mutant strains encoding varying lengths of the region upstream of OueS integrated at the native *ompU* locus were monitored for biofilm formation. All strains repressed biofilm formation similar to the ectopic OueS expressing clone (**Fig 1H**). An *ompU* promoter mutant strain (ΔP*_ompU_*) behaves like the Δ*ompU* strain indicating that OueS is expressed from the *ompU* promoter. Two-tailed unpaired T-test *p-*value: **** ≤ 0.00001.

**Figure S6. Variation within the OueS alleles.** *ompU* alleles were analyzed for sequence variation^41^ and OueS sequences were extracted from representative clades (A) Sequence alignment of the OueS loci from representative alleles demonstrates extensive variability in the region within the *ompU* ORF but is conserved in the 3’ UTR. (B) Secondary structure prediction reveals that the representative OueS alleles match one of the eight secondary structures. The structures requiring the least free energy (ΔG) are shown.

**Figure S7. Transcriptome of OueS alleles under virulence-inducing conditions**. Strains encoding a clinical-like allele (OueS^GBE1194^) and an environmental allele (OueS^GBE1116^) were selected based on mouse intestinal colonization data (**Fig 4B**) and transcriptomes were generated under virulence-inducing AKI conditions. Comparing the transcriptomes of two toxigenic-like and environmental alleles reveals that 96 and 66 genes are modulated uniquely by clinical-like and environmental OueS, respectively (**Table S9**).

### Supplementary Tables

Table S1. Transcriptomic changes during biofilm formation associated with *ompU* expression.

Table S2. Differentially expressed genes in biofilms in *ompU* and *rpoE* mutants.

Table S3: OueS-regulated genes during biofilm formation.

Table S4: ToxR-regulated genes during biofilm formation.

Table S5. ToxR-regulated genes under virulence-inducing conditions.

Table S6. Overlap between the ToxR and OueS-regulated genes under virulence-inducing conditions.

Table S7: OmpU and RpoE regulons under virulence-inducing conditions.

Table S8: Genes modulated by OueS under virulence-inducing conditions.

Table S9: Transcriptomes of clinical-like and environmental-like OueS alleles under virulence-inducing conditions.

Table S10. Strains, vectors and primers used in this study.

