## Supplementary Figures for "Modular small RNA drives the emergence of virulence traits and environmental trade-offs in *Vibrio cholerae*"

### Supplementary Figure 1

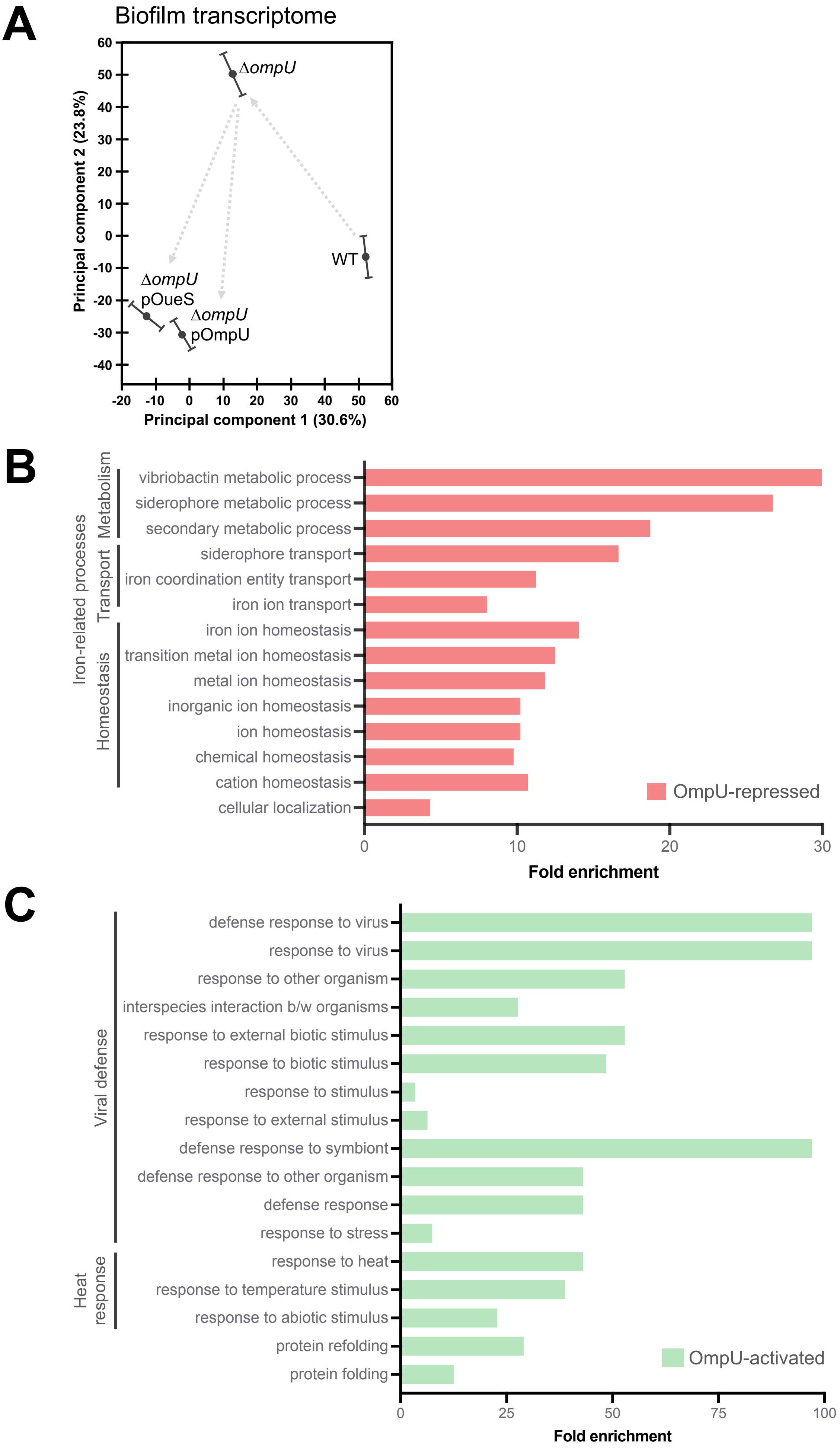

### Supplementary Figure 2

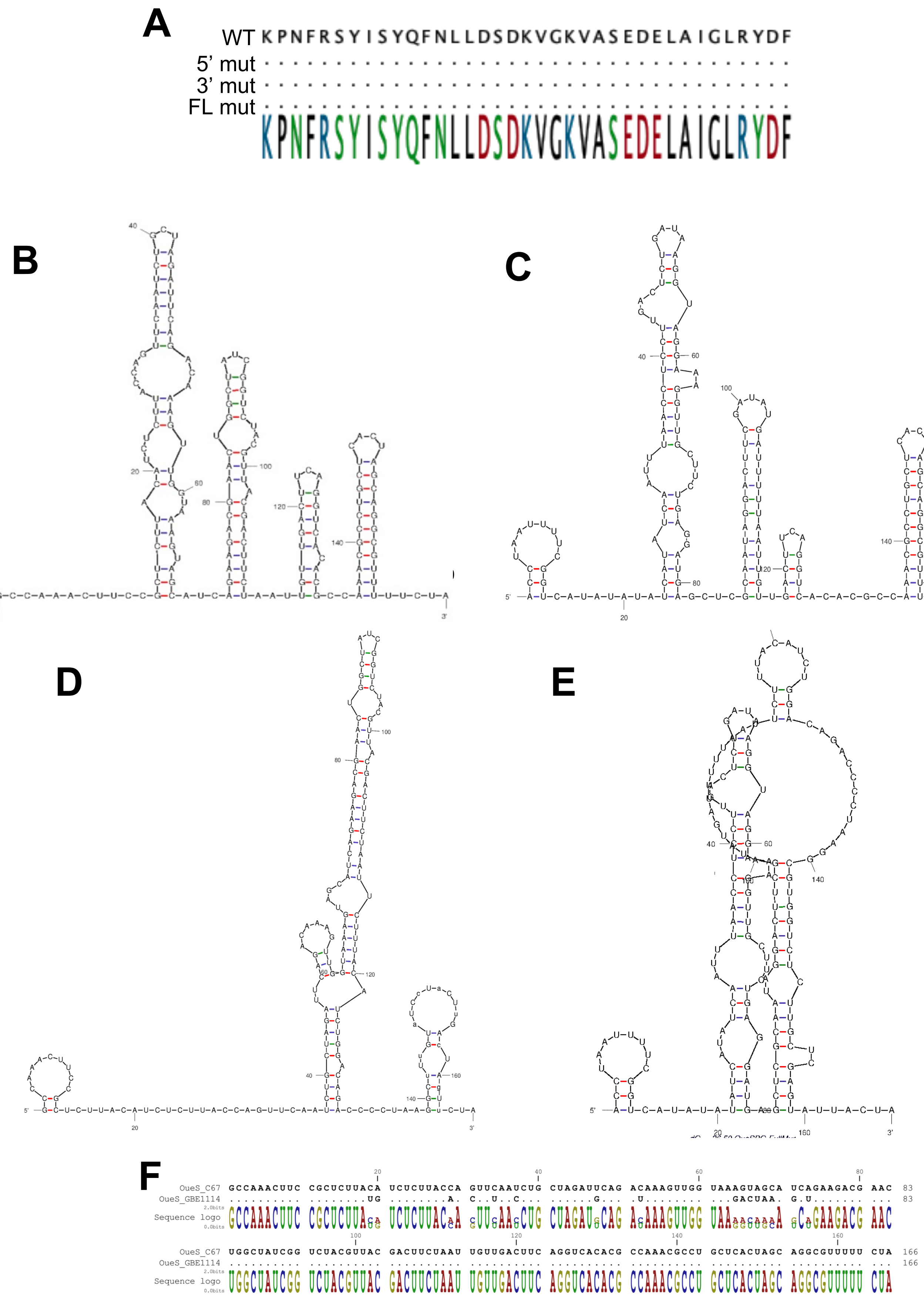

### Supplementary Figure 3

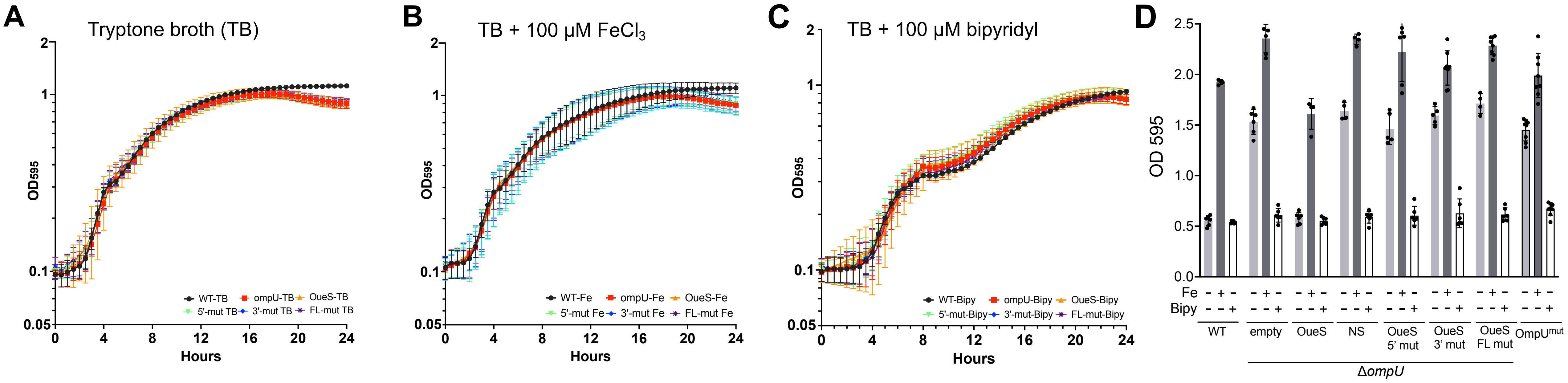

**Supplementary Figure 4**

**A**

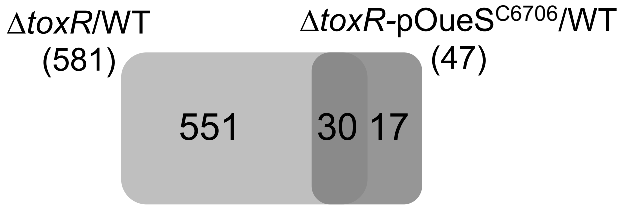


**B**

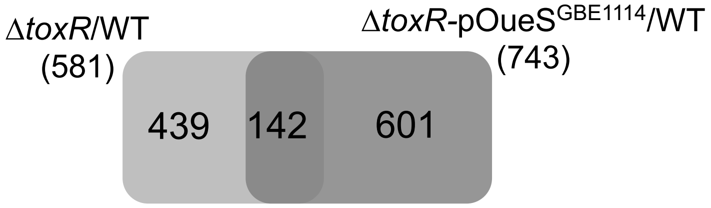

**Supplementary Figure 5**

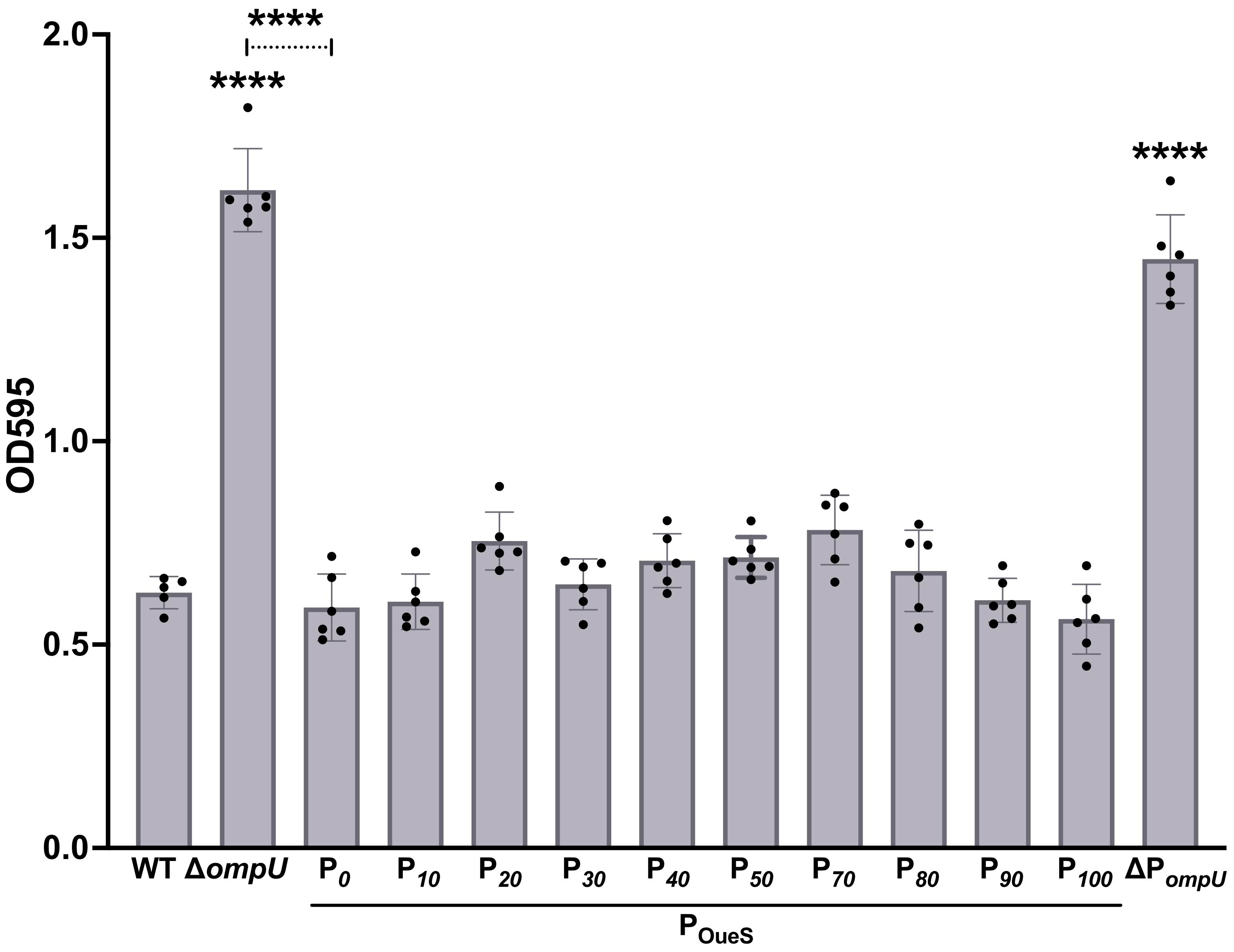

### Supplementary Figure 6

**A**

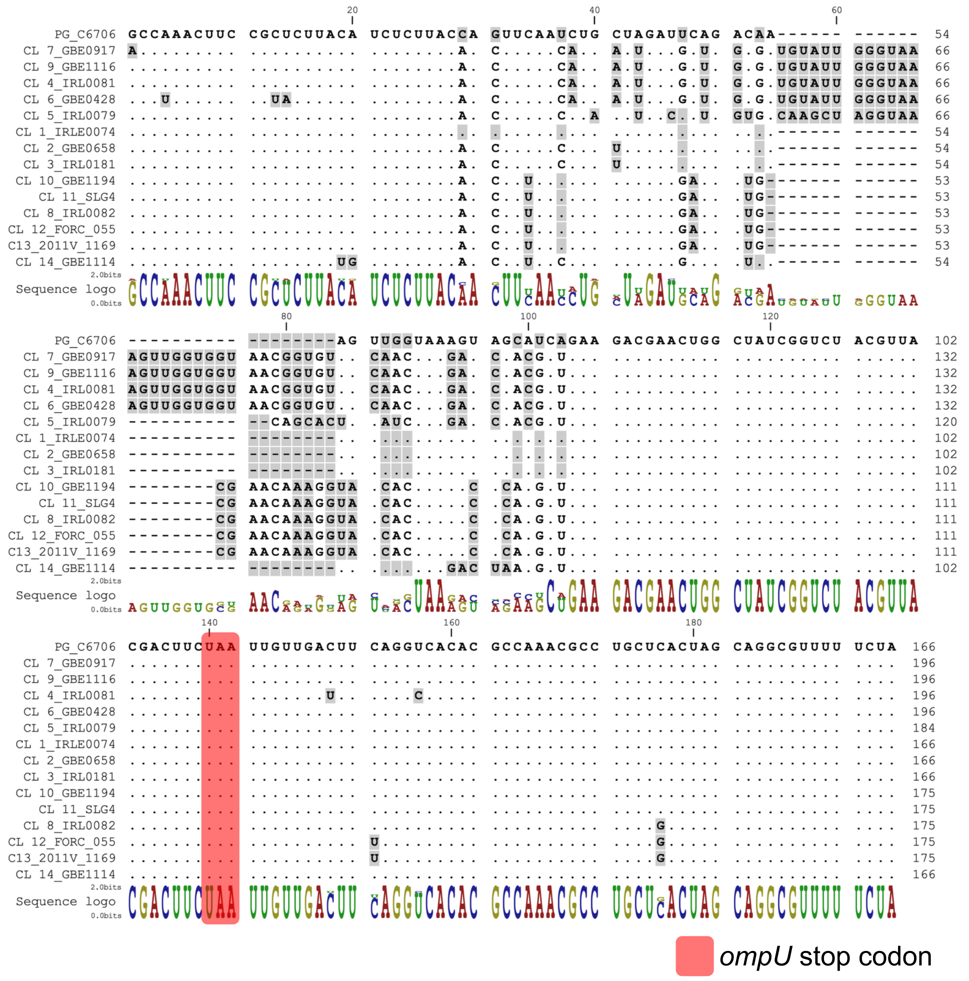


**B**

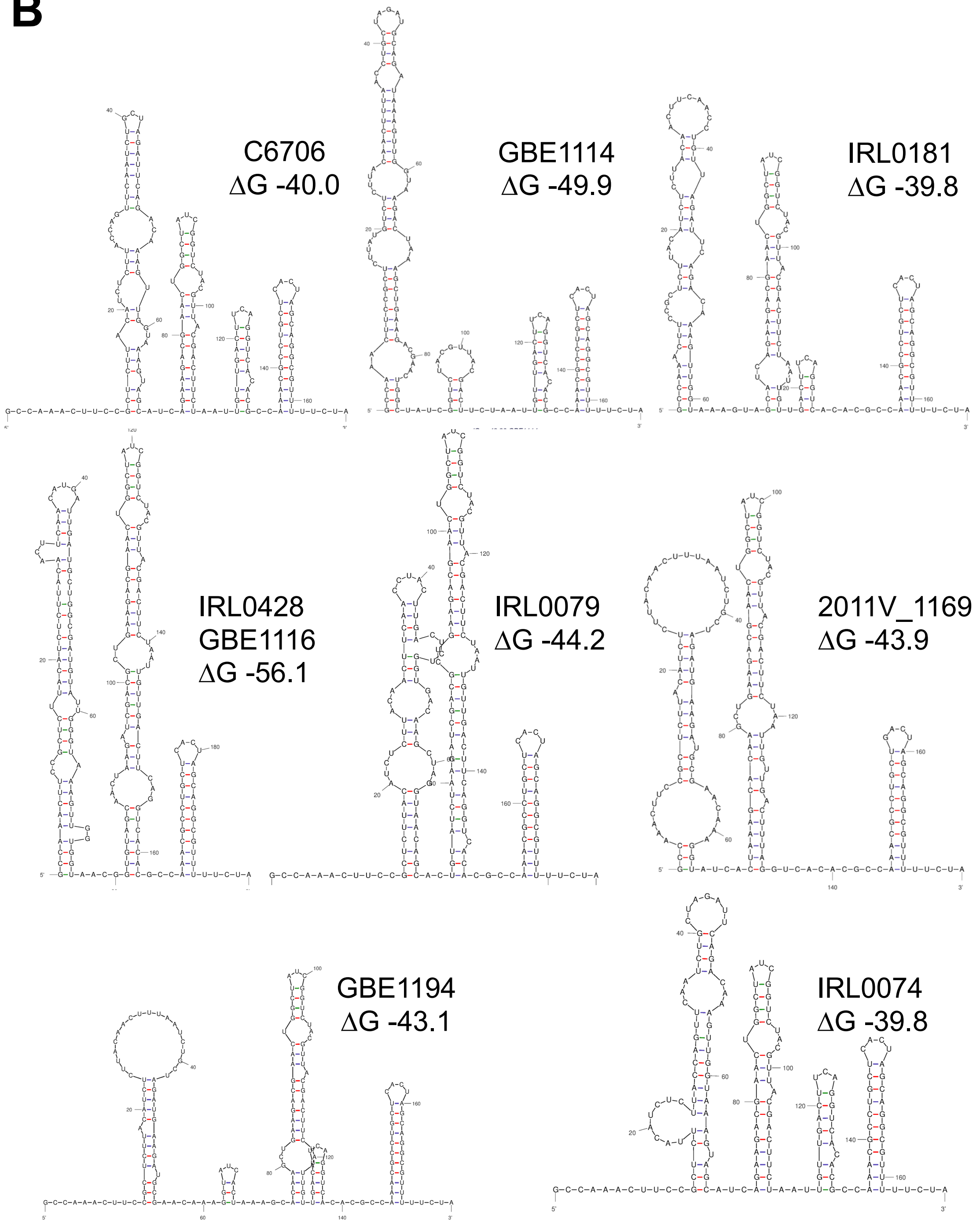

**Supplementary Figure 7**

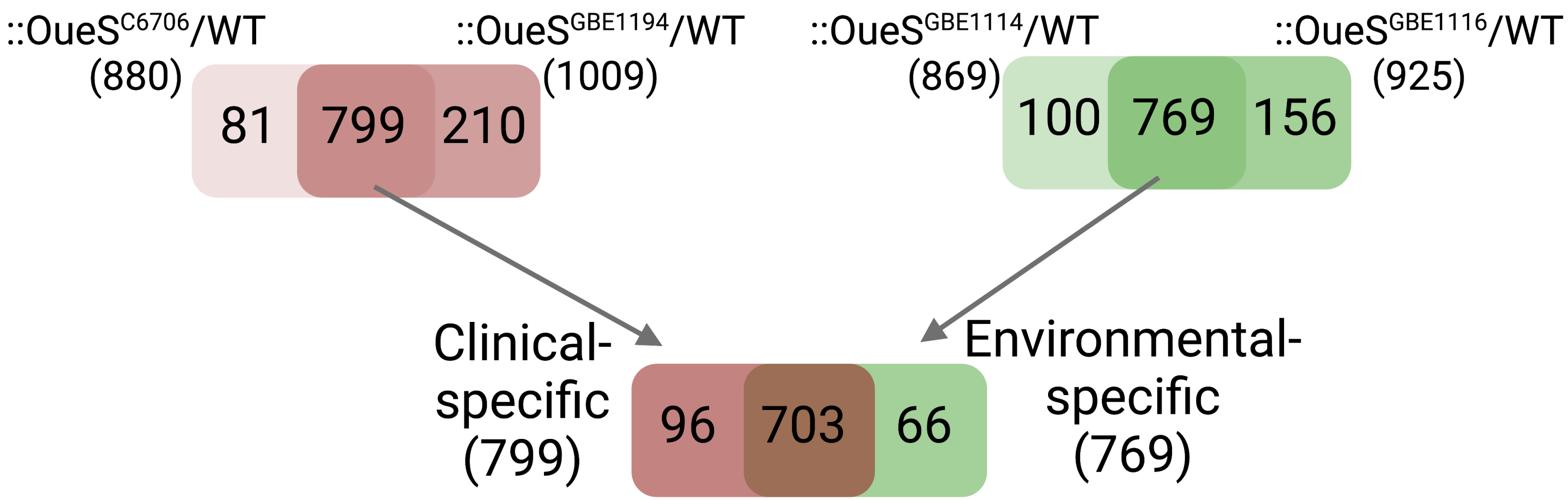
